# Loss of *Smad4* promotes aggressive lung cancer metastasis by de-repression of PAK3 via miRNA regulation

**DOI:** 10.1101/2021.04.29.441956

**Authors:** Xiaohong Tan, Lu Tong, Lin Li, Jinjin Xu, Shaofang Xie, Lei Ji, Jujiang Fu, Qingwu Liu, Shihui Shen, Yun Liu, Yanhui Xiao, Feiran Gao, Robb E. Moses, Nabeel Bardeesy, Yanxiao Wang, Jishuai Zhang, Kwok-kin Wong, Longying Tang, Lei Li, Dianwen Song, Xiao Yang, Jian Liu, Xiaotao Li

## Abstract

Over 85% of lung cancer patients harbor overt or subclinical metastases at diagnosis, and therefore most patients die of progressive metastatic disease despite aggressive local and systemic therapies. Somatic mutations in the *Smad4* gene have been found in non-small-cell lung cancer, but the underlying mechanism by which Smad4 loss-of-function (LOF) accelerates lung cancer metastasis is yet to be elucidated. Here, we generated a highly aggressive lung cancer mouse model bearing conditional *Kras^G12D^*, *p53^fl/fl^* LOF and/or *Smad4 ^fl/fl^* LOF mutations. The *Smad4^fl/fl^*; *p53 ^fl/fl^; Kras^G12D^* (SPK) mutant mice manifested a much higher incidence of tumor metastases than the *p53 ^fl/fl^; Kras^G12D^* (PK) mice. Molecularly, PAK3 was identified as a novel downstream effector of Smad4, mediating metastatic signal transduction via the PAK3-JNK-Jun pathway. Upregulation of PAK3 by Smad4 LOF in SPK mice was achieved by attenuating Smad4-dependent transcription of miR-495 and miR-543. These microRNAs (miRNAs) directly bind to the PAK3 3’UTR for blockade of PAK3 production, ultimately regulating lung cancer metastasis. An inverse correlation between Smad4 and PAK3 pathway components suggests clinical use of Smad4 LOF as a potential marker for prognosis in human lung cancer. Our study highlights the Smad4-PAK3 regulation as a point of potential therapy in metastatic lung cancer.

## Introduction

The American Cancer Society compiles on cancer incidence, mortality and deaths occurring in the United States every year. Lung cancer is the leading cause of cancer deaths worldwide, accounting for more solid tumor deaths than breast, pancreatic, prostate, and colorectal combined^1–5^. Lung cancer is broadly divided into small-cell lung cancer and non-small-cell lung cancer. More than 85% of lung cancers are classified as non–small-cell lung cancer (NSCLC) ^6^. Activating mutations of the *Kras* gene, found in 30 to 50% of NSCLC samples, are one of the most common genetic alterations in human lung cancer^7–9^. In addition, mutations of *Trp53* have been frequently reported in lung cancer (50-75%)^10^. Mutant *Kras* (here after called *Kras^G12D^*) alone can initiate lung cancer in mice, however, the tumors rarely metastasize ^11^. Cre/LoxP technology makes it possible to develop multiple conditional alleles of tumor suppressor genes or oncogenes and to initiate tumors with short latency and high penetrance ^12^. In conditional lung cancer models based on nasal delivery of adenoviral CRE (*adeno-Cre*), only a subset of cells acquires mutations within the lung, mimicking the sporadic tumorigenic process ^13^. Concomitant expression of *Kras^G12D^* ^14^ and *p53^fl/fl^* ^15^ in mouse models leads to a histologically and invasively more “humanized” version of NSCLC lung cancer^16^. Therefore, the *p53^fl/fl^*; *Kras^G12D^* mouse models realistically mimic the developmental stages of human lung cancer.

*Smad4*, a tumor suppressor, is the central intracellular mediator of TGF-β signaling. Smad4 inactivation is associated with different types of cancer. For example, loss of *SMAD4* is strongly associated with increased metastatic potential and promotes pancreatic and colorectal cancer progression^17^. Somatic mutations in the *Smad4* gene have also been described in NSCLC^18, 19^. Smad4 deficient lung metastases show a significant correlation with CCL15 expression based on patient specimens^20, 21^. The TGFβ-induced Smad4 complex stimulates the expression of SNAIL1 and TWIST1, which act as transcriptional factors repressing the expression of E-cadherin (EMT marker) ^22^. Depletion of Smad4 also resulted in a substantial upregulation of MAPK-JNK signaling pathways ^17, 23^. In animal models, *Smad4* deficiency blocks TGFβ-driven epithelial to mesenchymal transition in cancer progression through multiple factors ^14, 24^. However, the potential mechanism of Smad4 in lung cancer metastasis has not been elucidated *in vivo*.

In this study, we used *adeno-Cre* to conditionally activate a *Kras^G12D^* allele with concomitant deletion of *Smad4* (*Smad4 ^fl/fl^*) and *p53* (*p53^fl/fl^*) genes to induce lung cancer. Expression of mutant *Kras^G12D^* along with *p53* and *Smad4* loss-of-function (LOF) engendered a high incidence of metastasis to different tissues, compared to that found in *p53^fl/fl^*; *Kras^G12D^* mice. We found Smad4 deficiency promotes PAK3 elevation by attenuating expression of miR-495 and miR-543, inhibitory factors for PAK3. Furthermore, activation of the PAK3-JNK-Jun pathway in the *Smad4 ^fl/fl^*; *p53^fl/fl^*; *Kras^G12D^* triple-mutant mice contributed to the metastatic potential, suggesting a possible new target for therapy of non-small cell lung cancer. Action of Smad4 in the regulation of PAK3 offers a tool for lung cancer prognosis. This study provides novel insights into Smad4-dependent regulation of tumorigenesis, progression and metastasis in lung cancer.

## Results

### *Smad4* deletion accelerates lung tumorigenesis and metastasis

To investigate the contribution of Smad4 LOF in lung cancer progression and metastasis in the context of conditional mouse lung cancer models, we utilized existing Cre/LoxP–controlled, genetically engineered mouse models with Kras (*Kras^G12D^*) ^14^, p53 LOF (*p53^fl/fl^*) ^15^, and Smad4 LOF (*Smad4^fl/fl^*) ^24, 25^ mutations to generate a cohort of lung tumors by nasal delivery of adenovirus expressing Cre recombinase (adeno-Cre). Mice with a homozygous deletion of *Smad4* alone in lungs after adeno-Cre treatment survived beyond 52 weeks and didn’t develop any gross anatomic abnormalities or lung cancer (data are not shown). Mice with *Kras^G12D^* activation (Abbreviated as *K*) alone developed lung tumors following a long latency comparable to previous reports^26, 27^, with no detectable metastasis (Fig. 1A). Inactivation of *Smad4* in the background of Kras^G12D^ mutation (SK) had a metastasis frequency of 5.6% (Fig. 1A, S1A). However, tumors in *Smad4 ^fl/fl^*; *p53 ^fl/fl^; Kras^G12D^* (SPK) mutant mice had a dramatically higher metastatic rate (51.2%, Fig. 1A, S1A) than the traditional *p53 ^fl/fl^; Kras^G12D^* (PK) metastatic model (13.6%, Fig. 1A, S1A, and Sup. Table 1). The malignant manifestation included metastatic lesions to heart, thorax, and thymus, in addition to bone, kidney, liver, and peritoneum that are commonly observed in PK mice (Fig. 1B).

**Figure 1.**
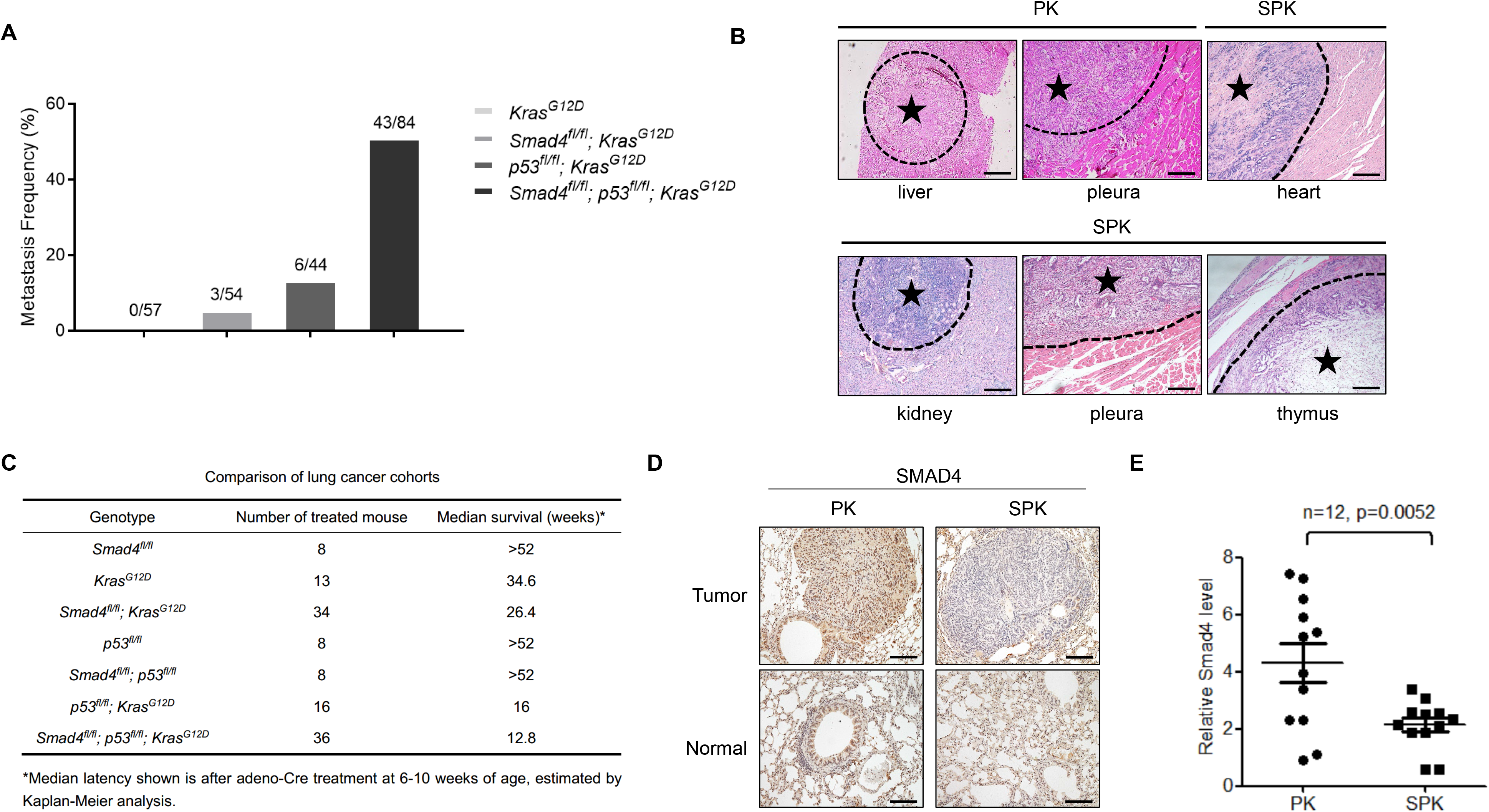
Smad4 deletion accelerates lung tumorigenesis and metastasis. **A.** Statistical analyses of lung cancer metastases frequency in *KRAS^G12D^* (n=57), *Smad4^fl/fl^*; *KRAS^G12D^* (n=54), p53^fl/fl^; *KRAS^G12D^* (n=44), *Smad4^fl/fl^*; *p53^fl/fl^*; *KRAS^G12D^* (n=84) mice. **B.** Representative images of H&E stained lung cancer metastatic tissues, including liver, pleura from PK mouse and heart, kidney, pleura, thymus from SPK mouse. Asterisk indicates the area of metastatic tumors. Scale bar, 100 μm (magnification, ×10). **C.** Survival statistics of different genotype mice with lung cancer. Median latency refers to the survival time on average (in weeks) after adeno-Cre treatment at the age of 6-10 weeks, estimated by Kaplan–Meier analysis. **D.** Representative IHC staining of Smad4 on normal and lung tumor sections of PK or SPK mice. Mice were all treated by adeno-Cre for 14 weeks. Scale bar, 25 μm (magnification, ×40). **E.** The Smad4 mRNA expression of PK and SPK mouse lung tumors was evaluated by real-time quantitative PCR. n=12; data represent means ± s.e.m.; as determined by Student’s *t* test.

Consistent with the metastatic rate in each group, the median survival duration for SPK, PK, SK, and K mice was 12.8, 16, 26.4, and 34.6 weeks respectively (Fig. 1C). Of note, Kaplan-Meier survival analysis showed that PK mice had prolonged median life-span compared to SPK mice (Fig. S1F). Most tumors exhibited features of lung cancer histologically and molecularly, as demonstrated by expression of TTF1 marker (Fig. S1B), with higher tumor burdens in SPK than in PK mice (Fig. S1C-1E). Meanwhile, we observed less than 20% sporadic lung tumors resembling a feature of squamous cell carcinoma, such as the typical nests of neoplastic squamous cells with positive staining p63 and Krt5 (Fig. S1G). Efficient ablation of Smad4 in SPK tumors was verified by immunohistochemical analysis of lung cancer tissues from SPK vs. PK mice (Fig. 1D) and by RT-PCR analysis of tumors for *Smad4* RNA expression (Fig. 1E). These data suggest that Smad4 suppresses tumorigenesis and metastasis of lung cancer in mouse model.

### Abrogation of *Smad4* promotes lung cancer cell migration and invasion

To address the action of Smad4 depletion in lung cancer cells with p53 LOF and Kras^G12D^ mutation in promoting lung cancer metastasis, we isolated primary lung cancer cells from PK and SPK mouse tumors and generated immortalized PK and SPK lung cancer cell lines (Fig. 2A). Transwell assays demonstrated that SPK cells were more invasive than PK cells (Fig. 2B, S2A). Furthermore, wound-healing assays substantiated an increased cell migration in SPK cells compared to PK cells (Fig. 2C, S2B).

**Figure 2.**
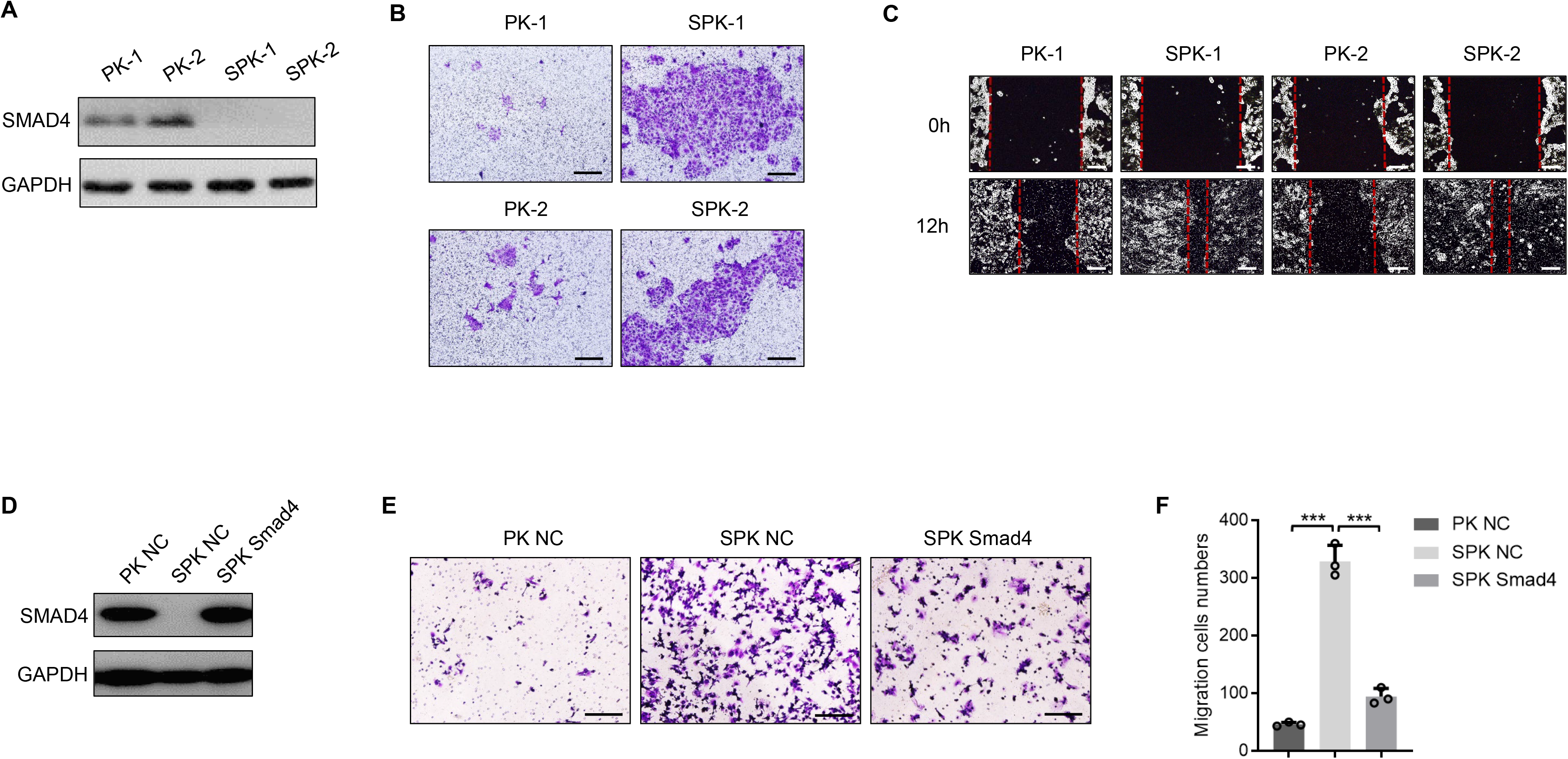
Abrogation of Smad4 promotes lung cancer cell migration and invasion. **A.** Primary lung cancer cells were isolated from lung tumors of PK and SPK mice and immortalized. The protein levels of Smad4 were examined in PK and SPK cells. Each lane represents a cell sample from an individual mouse. **B.** The role of Smad4 in murine lung cancer cell migration/invasion was tested by Transwell assay. **C.** Smad4 inhibited wound healing in cell culture. Cells were made a wound between the two red dashed lines, the area of the two dashed lines represent the level of wound healing and the cell migration activity. Wound healing percentage is the ratio of the wound healing area and the primary wound area. **D.** Transfection of Smad4 plasmid in SPK cells increased the protein level of Smad4. **E.** Overexpression of Smad4 in SPK cells decreased cell migration/invasion. **F.** The number of migrated cells was quantified. Data represent means ± s.e.m. ***, p<0.001, as determined by Student’s *t* test.

Actin assembly provides a major force for cell movement by driving lamellipodia and filopodia that propel the leading edge ^28^. To determine the impact of Smad4-depleted H1299 cells on the dynamic changes in lamellipodia and filopodia, stable knockdown of Smad4 by a specific shRNA (shSmad4) in the human lung cancer cell line H1299 (H1299-shSmad4, Fig. S2C), which has an activated *RAS* and p53 LOF mutations, stimulated by serum were evaluated by cell spreading and morphological changes (Fig. S2D). Indeed, F-actin staining revealed an enhanced formation of lamellipodia in cells silencing Smad4 (Fig. S2E). To determine how critical Smad4 is in SPK cell invasion, we performed a “rescue experiment” by reintroducing Smad4 into SPK cells (Fig. 2D). Re-expression of Smad4 significantly attenuated SPK cell migration and invasion in a transwell study (Fig. 2E, 2F), while 6 knocking down Smad4 enhanced invasion in human lung cancer cells (Fig S2F). Examination of the expression of EMT markers revealed that SNAIL1 was increased upon loss of Smad4 while no significant changes were observed in the expression of E-cadherin and TWIST1 between PK and SPK cells (Fig. S2G). Overall, these results demonstrate that silencing Smad4 in the context of p53 LOF and *Kras* mutation significantly promotes migration and invasion in human and murine lung cancer cells.

### PAK3 is a downstream effector of Smad4 mediating lung cancer cell metastasis

To understand the molecular mechanisms by which loss of function of *Smad4* promotes lung cancer cell metastasis, we compared gene expression profiles between SPK and PK model cells by RNA-sequencing (RNA-seq) analysis (Figure 3A) which disclosed 3,777 differentially expressed genes (DEGs) (Sup. Data 1), including an increased SNAIL1 mRNA expression. We then conducted a KEGG pathway analysis on these DEGs, showing cancer as the top enriched human disease (Figure 3B and Sup. Data 2). Moreover, the top enriched cellular processes included cell growth and death as well as cell motility (Figure 3B and Sup. Data 3). To identify metastasis-associated cancer genes, we focused on genes overlapped in cancer and cell motility categories (Figure 3C and Sup. Data 4). Among the cancer genes involving cell motility, PAK3 was the top DEG after ablation of *Smad4* (SPK *vs.* PK) (Figure 3C). We validated higher PAK3 protein and RNA expression in SPK lung tumors compared to PK tumors by immunohistochemical staining and quantitative PCR (Fig. 3D, 3E). Despite that Smad4 interacts with R-Smads to potentiate TGFβ signaling, our results demonstrated that regulation of PAK3 by Smad4 was not affected by the treatment of TGFβ or silencing Smad3 (Fig. S3A), suggesting a TGFβ-independent mechanism. To substantiate the function of PAK3 is indeed correlated with SMAD4, we generated stable knockdown of PAK3 in a SPK cell line (Fig. 3F). In wound healing assays, PAK3 knockdown clones (shPAK3 #1, shPAK3 #2) exhibited a reduction in wound closure compared with controls (shN) (Fig. 3G, 3H). Transwell assays demonstrated a much slower invasion of shPAK3 cells through the Matrigel than shN cells (Fig. S3B, S3C), suggesting that PAK3 is a downstream effector of SMAD4 to promote cell metastasis. To evaluate the regulation of PAK3 in cell metastasis, we performed gain-of-function experiments using constitutively active PAK3 stably integrated in H1299 cells. Cells expressing active PAK3 (H1299-caPAK3) migrated and invaded dramatically faster than cells with a vector control (H1299-vector) (Fig. S3D-3G). To assure the specific regulation of PAK3 by SMAD4, we measured the expression of PAKs (PAK1, PAK2, and PAK3) in PK and SPK cells with or without the treatment of TGFβ or TGFβ in the presence or absence of TGFβ inhibitor SB compound. We found that only PAK3, but not PAK1 or PAK2, mRNA levels are significantly upregulated upon loss of SMAD4 in a TGFβ-independent manner (Fig. S3H). The TRI gene, a target of TGFβ signaling, was included as a positive control (Fig. S3H). The regulation of PAK3 by SMAD4 in a TGFβ-independent manner may explain the partial changes in EMT markers. Our data suggest that the PAK3, a downstream effector of SMAD4, mediates lung cancer cell metastasis.

**Figure 3:**
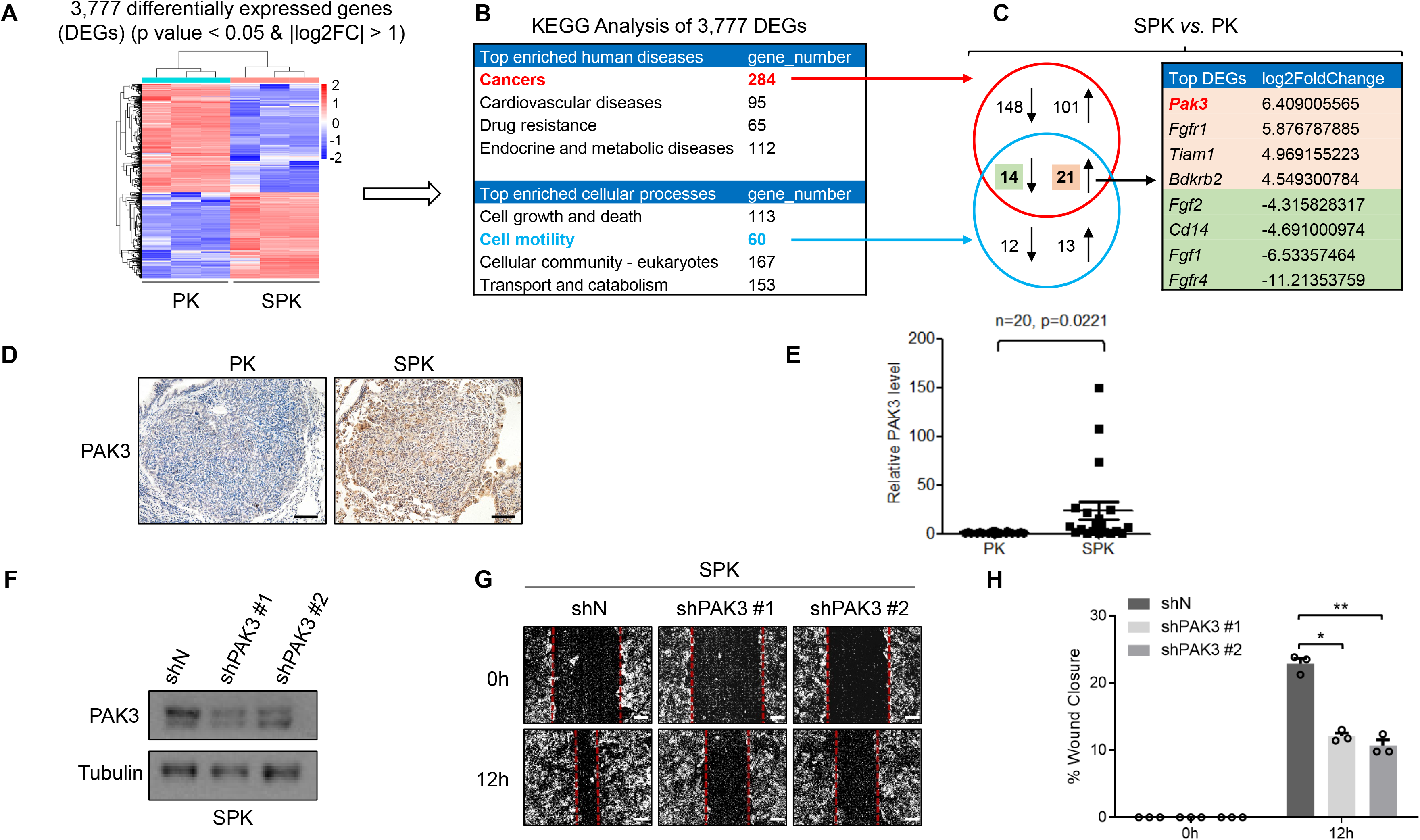
*PAK3* is a downstream effector of Smad4 mediating lung cancer cell metastasis. **A.** Gene expression profiles were detected between PK and SPK cells by RNA-sequencing(RNA-seq) analysis. **B.** KEGG Analysis of 3,777 DEGs **C.** Analysis of DEGs overlapped between cancer and cell motility KEGG pathways. **D.** Representative IHC images of PAK3 expression in PK and SPK mouse lung tumors. Scale bar, 50 μm (magnification, ×20). **E.** Quantitative analysis of *PAK3* RNA expression in PK and SPK mouse lung tumors. Each spot represents a sample from an individual mouse. n=20, data represent means ± s.e.m.; p<0.001, as determined by Student’s *t* test. **F.** Western blots showing decreased protein levels of PAK3 in shPAK3 (#1/#2) cells. **G.** The migration ability of SPK shN and shPAK3 (#1/#2) cells was tested by wound healing assay. Cells were made a wound between the two dashed lines, the area of the two dashed lines represents the level of wound healing and the cell migration activity. **H.** Statistical analyses of wound closure percentage. Data represent means ± s.e.m. *, p<0.05, **, p<0.01, as determined by Student’s *t* test.

### PAK3 enhances the JNK-Jun signal pathway

Given the inverse correlation between SMAD4 and PAK3 in PK/SPK lung cancer cells, we wondered how PAK3 signaling affects cell migration and invasion. In the PAK family proteins, PAK1 has been shown to promote cancer cell migration^29^. The c-Jun NH2-terminal kinase (JNK) is activated in PAK3 transfected cells, and inhibition of JNK activity abolishes PAK3-mediated cell migration in neuroendocrine tumors^30^. We also validated that blocking JNK activity abolished the motility difference between PK and SPK cells (Fig. S4H), suggesting that JNK mediates PAK3 signaling. To determine whether PAK3 may regulate JNK-Jun activities in lung cancer cells, we checked the expression of p-JNK and p-Jun in PK and SPK cells. Lack of Smad4 and elevation of PAK3 in SPK induced significant phosphorylation of JNK with Jun (p-Jun) activation (Fig. 4A), as well as higher c-Jun mRNA levels in SPK cells than in PK cells (Fig. S4G). Similar activation of the JNK-Jun pathway was observed in the Smad4-deficient H1299-shSmad4 cells (Fig. S4A). In contrast, re-expression of exogenous Smad4 in SPK cells significantly suppressed the level of PAK3 and activation of the JNK-Jun kinases (Fig. 4B), suggesting that inhibition of Smad4 promotes a PAK3-dependent activation of JNK and Jun. In line with positive regulation of PAK3 on the JNK-Jun kinases, we detected enhanced level of P-MEK (S298), a direct downstream target of the PAK3, in SPK cells compared with PK cells (Fig. S4I). These data demonstrate that PAK3 is activated after loss of Smad4.

**Figure 4:**
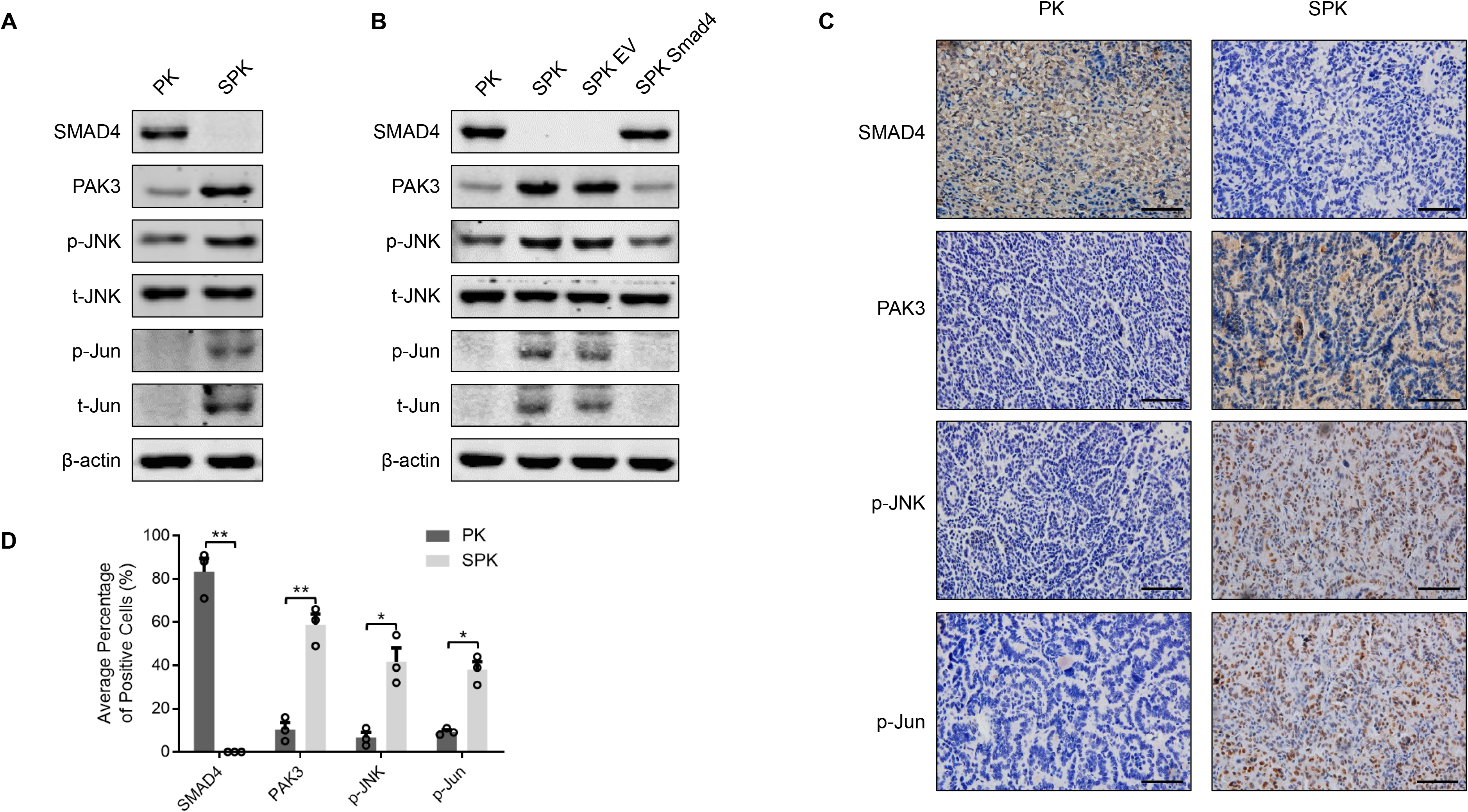
PAK3 enhances the JNK-Jun signal pathway. **A.** PK and SPK cells were analyzed for the expression of Smad4, PAK3 and JNK-Jun signaling related factors by Western blotting. **B.** Western blots showing the effects of Smad4 overexpression in SPK cells. **C.** IHC staining detected the expression of Smad4, PAK3, p-JNK and p-Jun in mouse lung tumors of PK and SPK mice. Scale bar, 25 μm (magnification, ×40). **D.** Quantification of average percentage of Smad4, PAK3, p-JNK and p-Jun positive cells in mouse lung tumors of PK and SPK mice. Data represent means ± s.e.m. *, p<0.05; **, p<0.01, as determined by Student’s *t* test.

To substantiate the role of PAK3 in regulating the JNK pathway, we silenced PAK3 in H1299 cells with a panel of shRNAs and found a drastic reduction in p-JNK and p-Jun levels (Fig. S4B). Conversely, the levels of p-JNK and p-Jun were significantly increased in H1299 cells expressing a constitutively active PAK3 (caPAK3, Fig S4C), suggesting that elevated PAK3 indeed activates the JNK-Jun signal pathway. Immunohistochemical analyses substantiated a positive staining of PAK3, p-JNK, and p-Jun that are negatively correlated with Smad4 in SPK lung tumor tissues compared to the PK samples (Fig. 4C, 4D). Therefore, our data suggest that PAK3 acts in signal transduction between Smad4 and the JNK-Jun signal pathway in lung cancer cells.

### SMAD4 negatively regulates PAK3 via transactivation of miR-495 and miR-543 expression

As a transcription factor, SMAD4 may directly regulate PAK3 expression. Therefore, we performed bioinformatics analysis of the SMAD4 ChIP-Seq data from both human and mouse lung cells^31, 32^. However, we did not observe SMAD4 binding on the PAK3 promoter region (Fig. S4D and S4E). Neither did we find that exogenous expression of SMAD4, SMAD3, or SMAD2 could affect PAK3 promoter activity in PAK3 promoter-luciferase assays (Figure S4F). We then turned to the possibility of indirect regulation via microRNAs (miRNAs). To determine if SMAD4 regulates *PAK3* expression through its 3′ UTR, a known target region of miRNAs, we constructed a luciferase reporter transcriptionally fused *PAK3* 3′ UTRs down-stream of the firefly luciferase gene (Fig. 5A). Interestingly, transfection of the *PAK3* 3′ UTRs luciferase reporter resulted in much higher luciferase activity in SPK than in PK cells (Fig. 5B). However, transfection of SMAD4 into SPK or H1299 cells dramatically reduced the luciferase activities (Fig. 5C, and S5A-C), leaving a potential regulatory link between SMAD4 and PAK3-targeting miRNAs. With bioinformatics analysis, we predicted that 7 miRNAs may target both homo PAK3 and Mus PAK3 3′ UTRs (Fig.S5D, indicated by stars). Screening analysis indicated that three miRNAs, miR-495, miR-539 and miR-543, were positively regulated by SMAD4 in a dose-dependent manner in H1299 cells under TGFβ treatment (Fig. S5E). Despite the high similarity between the homo and Mus PAK3 3′-UTR sequences (Fig. S5G), only miR-495 and miR-543 expressions were upregulated by overexpression of SMAD4 in SPK cells (Fig. 5D). Therefore, our subsequent studies focused mainly on miR-495 and miR-543.

**Figure 5.**
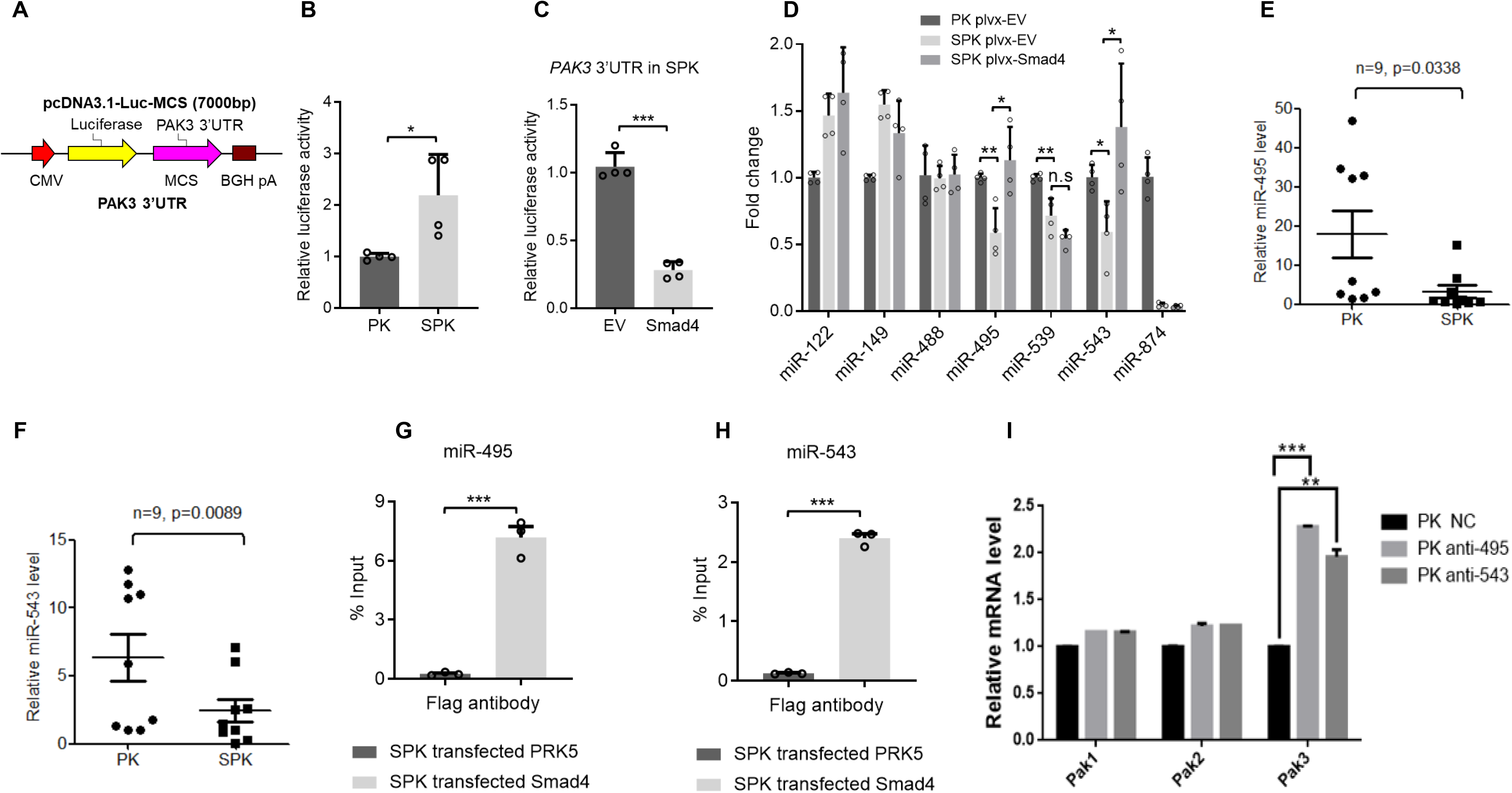
Smad4 negatively regulates PAK3 via transactivation of miR-495 and miR-543 expression. **A.** A luciferase reporter fused to PAK3 3′ UTRs was constructed in pcDNA3.1-Luc-MCS plasmid. **B.** Luciferase activities of PK and SPK cells transfected with PAK3 3’UTR-Luc plasmid confirmed increased PAK3 expression level SPK cells. Data represent means ± s.e.m. *p<0.05, as determined by Student’s t test. **C.** Luciferase reporter assay showing that overexpression of Smad4 decreased PAK3 expression level in SPK cells. Data represent means ± s.e.m. ***p<0.001, as determined by Student’s t test. **D.** RT-PCR detected expression of PAK3 correlated miRNAs in PK, SPK and SPK overexpressing Smad4 cells. Data represent means ± s.e.m. *p<0.05, **p<0.01, as determined by Student’s t test. **E-F.** Expression levels of miR-495 and miR-543 were determined in PK and SPK mouse lung tumors. n=9, data represent means ± s.e.m.; p<0.05, as determined by Student’s *t* test. G-H. The interaction between Smad4 and miR-495 (G) or miR-543 (H) in SPK cells were verified by ChIP assay. Data represent means ± s.e.m. **p<0.01, ***p<0.001, as determined by Student’s t test. **I.** qRT-PCR analysis gene expression in PK cells.

Consistent with the above findings, miR-495 and miR-543 levels were lower in SPK tumor samples (Fig. 5E, 5F). MiR-495 and miR-543 were decreased when Smad4 was transiently knocked down in H1299 cells (Fig.S6A-C). On the contrary, these two miRNAs were increased upon overexpression of Smad4 in H1299 cells (Fig. S6D-F). ChIP-qPCR analysis showed that SMAD4 bound to the promoters of miR-495 and miR-543 (Fig. 5G-H, and S6G-H). Moreover, transfection of antagomir miR495/miR543 in PK cells exclusively led to up-regulation of PAK3 mRNA without any effect on the expression of PAK1/2 (Fig. S5F). Therefore, our data indicate that SMAD4 affects PAK3 levels by positively regulating the expression of miR-495 and miR-543.

### MiR-495 and miR-543 directly bind to the PAK3 3’UTR and attenuate the metastatic potential of lung cancer cells *in vitro* and *in vivo*

To investigate whether miR-495 and miR-543 repress endogenous PAK3 expression, SPK cells were transfected with oligonucleotide mimics of miR-495 or miR-543. The expression of PAK3 was significantly reduced at both mRNA and protein levels following the treatment with miR-495 or miR-543 mimics (Fig. 6A, 6B). Similar inhibitory effects of these two oligonucleotides on PAK3 expression were obtained in H1299 cells (Fig. S7A, S7B). In contrast, transfected miR-495 or miR-543 inhibitors (antagomiR-495 or antagomiR-543) enhanced expression of PAK3 in both PK and H1299 cells (Fig. 6C, S7C). To determine if miR-495 and miR-543 act by directly targeting specific regions in PAK3 UTR, we generated mutant luciferase reporters with altered binding motifs of miR-495 and miR-543 in *PAK3* 3′ UTRs (Fig. S7D). Overexpression of miR-495 or miR-543 dramatically decreased the activity of the luciferase reporter bearing the WT 3′ UTR of *PAK3* (Fig. 6D, 6E). However, neither miR-495 nor miR-543 inhibited the activity of the mutant luciferase reporters (Fig. 6D, 6E), suggesting that miR-495 and miR-543 specifically bind to the PAK3 UTR.

**Figure 6.**
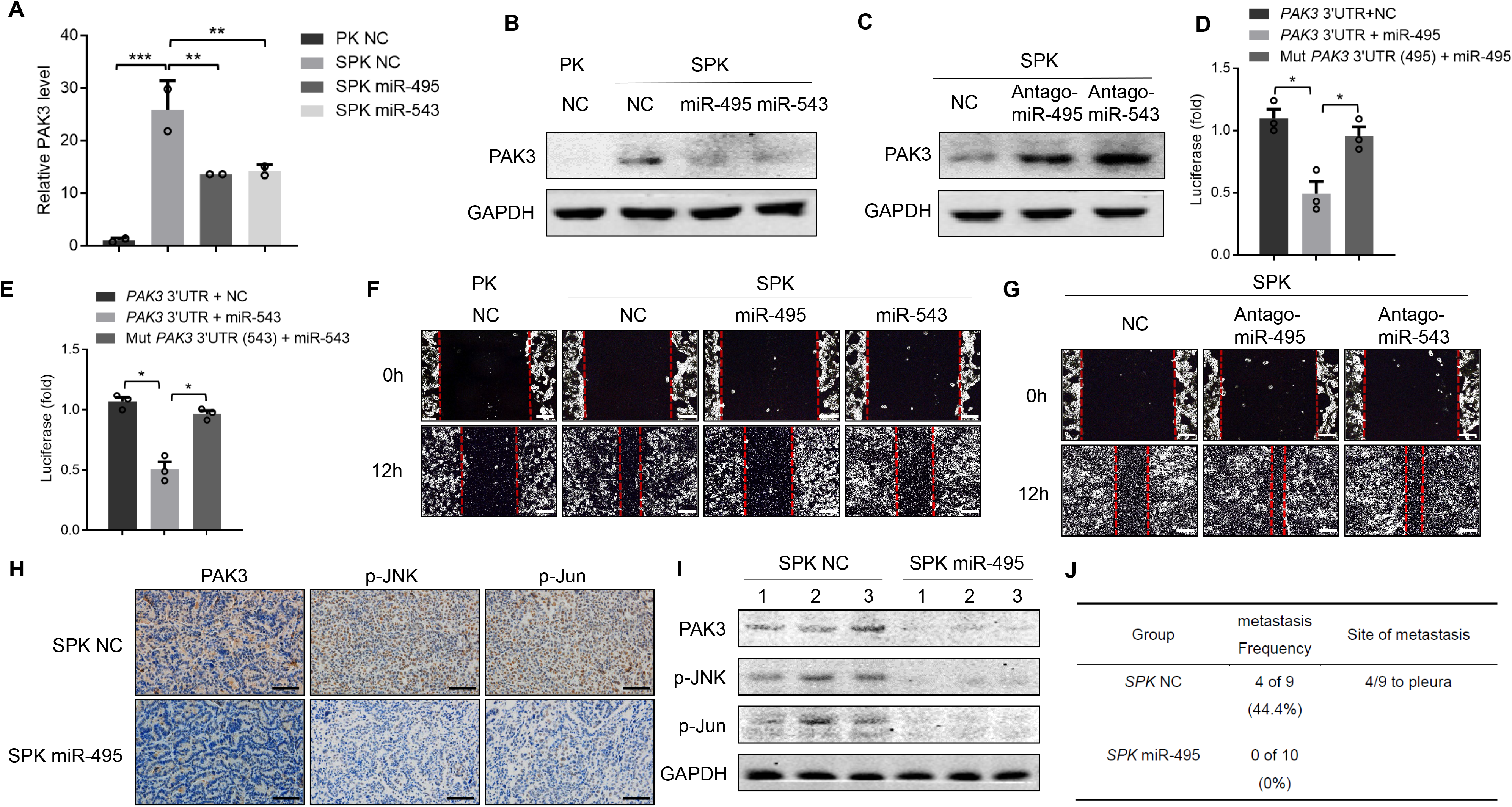
MiR-495 and miR-543 directly bind to the *PAK3* 3’ UTR and attenuate metastatic potential of lung cancer cells *in vitro* and *in vivo*. **A-B.** The expression of PAK3 was detected at both mRNA and protein levels after SPK cells were transfected with miR-495 or miR-543 mimics. **C.** The expression of PAK3 in SPK cells were detected following the transfection with miR-495 or miR-543 inhibitors (antagomiR-495 or antagomiR-543). **D-E.** PAK3 3’UTR and its mutation luciferase report system were constructed. The activity of the luciferase reporter bearing the WT 3’ UTR of *PAK3* was measured in H1299 cells transfected with miR-495 or miR-543 mimics. Luciferase activity means the expression level of *PAK3* 3’ UTR. Data are mean ± s.e.m. * p<0.05, as determined by Student’s *t*-test. **F.** Cells were made a wound between the two red dashed lines, the area of the two red dashed lines represent the level of wound healing and the cell migration activity. Cell migration ability was detected by wound healing assay in PK, SPK, SPK miR-495 and SPK miR-543 cells. **G.** Cell migration ability was detected by wound healing assay after antagomiR-495 or antagomiR-543 was introduced into SPK cells. **H-J.** 6-8 weeks SPK mice were treated by adeno-Cre and then injected NC agomir or miR-495 agomir after 4 weeks, once every two weeks. After 8 weeks, these mice were sacrificed to check lung cancer metastasis (**J**) and collected lung tumors to test the expression level of PAK3, p-JNK and p-Jun by IHC (**H**) and Western blotting (**I**). Scale bar, 25 μm (magnification, ×40).

Then, we assessed the impact of miR-495 and miR-543 on SPK/H1299 cell migration. MiR-495 or miR-543 mimics reduced PAK3 expression and attenuated wound closure in SPK cells (Fig. 6F, S7E) and H1299 (Fig. S7G). An increase in cell migration/invasion was observed when antagomiR-495 or antagomiR-543 was introduced into SPK cells (Fig. 6G, S7F) or H1299 cells (Fig. S7H). To determine if we could block or mimic the actions of miR-495/miR-543 with synthetic antagomirs or agomirs in mouse lung tumors, we tested miR-495/miR-543 mimics and antagomiR-495/antagomiR-543 in both male and female SPK/PK mice. To our surprise, only the miR-495 agomir can alter PAK3 expression *in vivo* (Fig. 6H, 6I), displaying downregulation of PAK3, p-JNK and p-Jun in mouse lung tumors. In the control group, SPK mice injected with miR-NC (control) had 44.4% metastasis, in which PAK3, p-JNK, and p-Jun activities were high (Fig. 6H-J, S8A-B). However, SPK mice injected with miR-495 mimics resulted in no tumor metastasis accompanied by reduced PAK3, p-JNK, and p-Jun levels in tumors (Fig. 6H-J). Our results demonstrate that the upregulation of PAK3 by *Smad4* LOF in SPK mice was achieved by attenuating SMAD4-dependent transcription of miRNAs that negatively regulate PAK3 expression, ultimately enhancing lung cancer metastasis.

### Correlation between SMAD4 and PAK3/JNK/Jun expression in human lung cancer samples

Given the importance of the SMAD4-PAK3-JNK-Jun axis in experimental lung cancer metastasis, we explored the possible clinical significance of SMAD4, PAK3, p-JNK, and p-Jun expression in human lung cancers, including 15 early, 12 advanced, and 30 metastatic human lung cancer samples along with 15 normal controls (Sup. Table 4). We found that SMAD4 was highly expressed in the controls, indicated by the highest protein levels (+++), while its expression continually dropped during the progression from early tumors to metastatic cancers. In contrast, the protein levels of PAK3, p-JNK, and p-JUN were increased in primary tumors and further in metastatic ones, compared with controls (Fig. 7A and 7B). By Pearson’s correlation analysis on a lesion-by-lesion basis, we found a strong negative correlation between SMAD4 and PAK3 or P-JNK or P-Jun with R values of −0.29, −0.77, and −0.73, respectively (Fig. 7C, S10A-C). Moreover, there were highly positive correlation R values between PAK3 and P-JNK or P-Jun (Fig. 7C, S10D-E). The above samples were also assayed for the expression of RAS (G12D) and P53 by immunohistochemistry along with organizational analysis by hematoxylin-eosin staining. The results showed that some of the metastatic samples had higher expression of RAS (G12D) and P53 (Fig. S9A-B), reflecting enhanced RAS activity and accumulation of mutantP53 in late stage of cancers^33, 34^. Consistent with our experimental findings, bioinformatics analysis of a published human cancer dataset (between 7 metastatic samples from lung cancers and 123 primary site lung tumor samples) ^35^demonstrated an overall reduction in *SMAD4* expression and an elevation of *PAK3* expression (Fig. S9C) displaying a strong negative correlation (Fig. S9D). These results suggested that reduced SMAD4 expression might be associated with poor prognosis in certain lung cancers. Taken together, we conclude that Smad4-mediated PAK3-JNK-Jun activation via regulation of miRNA in lung cancers appears to be a novel mechanism in the development of metastatic human lung cancers (Fig. 7D).

**Figure 7.**
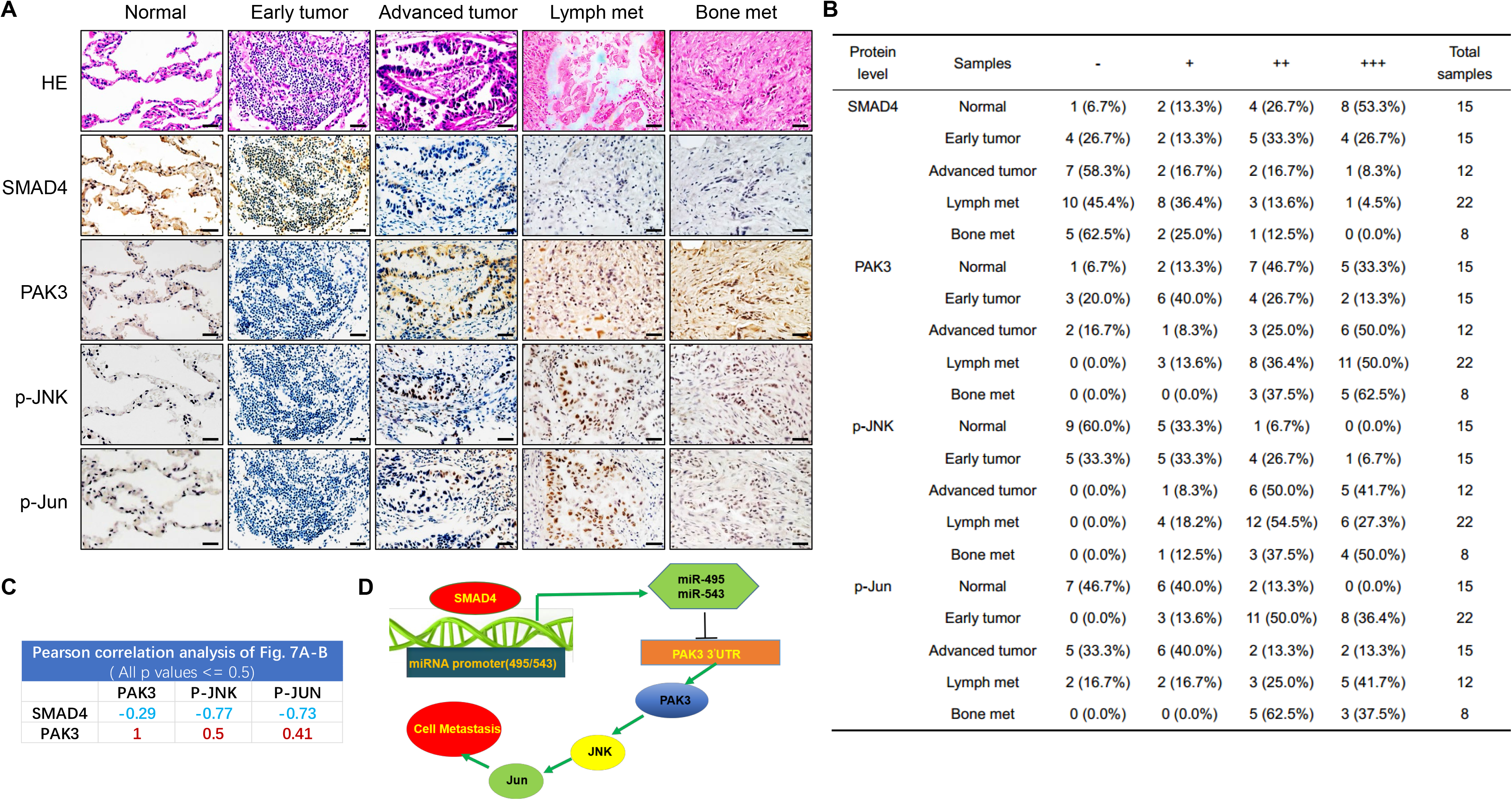
Correlation between Smad4 and PAK3/JNK/Jun expression in human lung cancer samples. **A.** Representative IHC images displaying an inverse correlation between Smad4, PAK3, p-JNK and p-Jun in human normal lung tissues, lung tumors and lung cancer metastatic samples. Scale bar, 25 μm (magnification, ×40). **B.** Statistics of positively stained percentages in human normal lung tissues (n = 15) and lung cancer metastatic tissues: early tumor (n = 15); advanced tumor (n =12); lymph nodes metastases (n = 22); bone metastases (n = 8). **C.** Pearson correlation analysis of protein expressions in Figure 7A-B. **D.** A proposed model for the role of Smad4 in lung cancer metastasis.

## Discussion

In our study, we have demonstrated that murine lung cancers with conditional *Smad4^fl/fl^*;*p53^fl/fl^*; *Kras^G12D^* mutations are a very aggressive metastatic model with a short survival duration after *adeno-Cre* delivery. We have validated the role of Smad4 LOF in promoting invasive and metastatic progression of lung cancer *in vitro* and *in vivo*. Our results implicate the Smad4-PAK3 signal events in metastatic progression of clinical lung cancer patients.

Smad4, the central mediator of TGF-β signaling, controls the signal transduction from cell membrane to nucleus, and has functions in many cellular processes, including proliferation, apoptosis and migration^36, 37^. As it is deleted in most pancreatic cancer, it is also called DPC4 (deleted in pancreatic cancer)^38^. Mutations of Smad4 have been detected in pancreas cancer, colon cancer, cholangiocarcinoma cancer and gastric cancers, suggesting an important tumor suppressor function of Smad4^39–43^. Smad4 heterozygous mice developed gastric cancer because of haploinsufficiency ^44^, therefore, specific Cre recombinase strategies were used to study the role of Smad4 loss in cancer development. Tissue-specific knockout of Smad4 could cause tumor formation in mammary tissue ^45^, including skin ^46^, liver ^47^, and colon ^48^. For lung tumor, TCGA data demonstrate heterozygous Smad4 loss in 13% of lung squamous cell carcinomas and 47% of lung adenocarcinomas^49, 50^. We have observed an association of reduced Smad4 expression with lung cancer malignancy in clinical metastatic samples, substantiating the role of Smad4 regulation in lung cancer progression.

Kras, p53 and Smad4 alterations are also frequently observed in other metastatic cancers, for example, in patients with metastatic colorectal cancer (CRC) with 38, 60 and 27% mutation rate respectively ^51^. CRC patients with a *Kras* mutation, *p53* mutation, or *Smad4* mutation, were at a higher risk of distant metastasis^52^. A number of classical pathways show crosstalk with the *Smad4* tumor suppressor, possibly explaining why *Smad4* LOF accelerates lung cancer metastasis in combination with *Kras* and *p53* mutation. A convergence of *p53* and Smad signaling pathway has been established^53^. Considering that some human lung tumors have not only SMAD4 mutation but also triple mutations in *KRAS*, *TP53*, and *SMAD4* (Sup. Tables 2-3)^54^, our findings may address the impact of *p53* mutation in *Smad4* LOF on the progression of these lung cancers. In addition, *Smad4* represents a barrier in *Kras*-mediated malignant transformation in a pancreatic cancer model^25, 55^. In this study, our finding that Smad4 inactivates PAK3-JNK-Jun pathway in advanced or metastatic lung cancer provides a new function for the tumor suppressive Smad4.

Several studies have indicated that TGF-β/SMAD4 signaling acts in regulation of different miRNAs. It has been shown that TGF-β1 induces miR-574-3p transcription to inhibit cell proliferation via SMAD4 binding to the promoter of miR-574-3p^56^. In addition, miR-23a-27a-24 cluster is induced by TGF-β1 in a Smad4-dependent manner ^57^. On the other hand, miR-19b-3p promotes proliferation of colon cancer cells by binding the 3′-untranslated regions (UTR) of *Smad4* directly ^58^. In this study, we have demonstrated for the first time that miRNAs miR-495 and miR-543 are transcription targets of *Smad4*.

In summary, this study identifies a new effector of Smad4, *PAK3*, and a novel miRNA-mediated mechanism in the regulation of lung cancer metastasis. We conclude that downregulation of *Smad4* de-represses the PAK3-JNK-Jun pathway via attenuated production of miR-495/miR-543 in lung cancers. Combination of experimental, clinical and bioinformatics analyses has demonstrated Smad4-PAK3 regulation acts in metastatic lung cancer malignancy. Determination of Smad4/PAK3 status may be of value in stratifying patients into treatment regimens related to personalized therapy.

## Materials and methods

### Experimental mice

To obtain the cohorts in our study, floxed *Smad4* allele ^24, 59, 60^, *Kras^G12D^* allele ^27^ and *p53* conditional allele (hereafter called *p53^fl/fl^*) ^15^ were used. PCR analyses of Kras and p53 allelic recombination were described in previous study^61^. Smad4 primer sets were used to confirm exon deletion. The primers were listed below: Kras-1: gtctttccccagcacagt; Kras-2: ctcttgcctacgccaccag; Kras-3: agctagccaccatggcttgagt; p53-1: cgcaatcctttattctgt; p53-2: agcacataggaggcaga; p53-3: tgagacagggtcttgct; Smad4-1: tgcattccagcctccca; Smad4-2: gccagcagcagcagacagac. All 6-8 weeks old PK and SPK mouse were treated with 1×10^6^ pfu Ad-Cre per mice by nasal drip to induce lung cancer. All experiments were conducted under specific pathogen free (SPF) conditions and handled according to the ethical and scientific standards by the Animal Center at Shanghai Key Laboratory of Regulatory Biology, Institute of Biomedical Sciences, School of Life Sciences, East China Normal University following procedures approved by the Institutional Animal Care and Use Committee. All strains were B6. 129. To evaluate the production of metastasis in mouse models, half of the organ sample was sectioned into slides (0.5 μm thickness per slide), and five representative slides were selected from each sample for the H&E staining. Once there are tumors found in organs other than lungs, immunohistochemical staining on TTF1 (a marker to demonstrate the origin of metastatic lesions from lung adenocarcinoma) would be conducted, and the tumors with the positive staining of TTF1 are considered as metastasized from the lungs.

### RNA extraction and RNA-seq

Total RNA of SPK and SPK model cells was extracted using RNeasy kit (Qiagen, Valencia, CA) and quantified with NanoDrop 1000 (Thermo Fisher Scientific, Waltham, MA). The cDNA sequencing libraries were prepared using Illumina’s TruSeq Sample Preparation Kit (San Diego, CA) and the sequencing was performed using Illumina Genome Analyzer. RNA-seq data were analyzed using R with various packages. The differential analysis of genes was conducted on counts using DESeq2 package. Differentially expressed genes (DEGs) were identified as such if the fold change > 2 and the p-value < 0.05. Gene ontology (GO) enrichment and enriched KEGG (Kyoto Encyclopedia of Genes and Genomes) pathways were performed.

### Cell culture

The mouse lung cancer derived cell lines PK and SPK were established from in vivo PK and SPK tumors. Briefly, tumors were dissected 12 weeks post Ad-Cre virus treatment, minced into small pieces and digested with collagenase for 1 hour at 37°C. Digested tissue were filtered through a 100 um filter, then a 40 um filter using excess cold PBS to wash cell through filter. Finally, the tumor cells were cultured in RPMI-1640 (Hyclone) with 10% FBS. The medium was changed every day until cells outgrew and stable immortalized cell lines were formed.

The human lung cancer derived cell lines H1299 were maintained in RPMI 1640 Medium (Invitrogen) with 10% fetal bovine serum (HyClone). shN (target sequence AGCGGACTAAGTCCATTGC) and shSmad4 (target sequence GGATTTCCTCATGTGATCT) plasmids in H1299 were kindly provided by Dr. Xinhua Feng. shN and shSmad4 were transfected to H1299 cells and selection at 2 ug/ml puromycin. The pGIPZ-PAK3 (target TAGTGCTTCGTTTACTTTG), The pGIPZ-PAK3 (target TGTATGCTCTGGTCTTGGT) were

obtained from Darmacon. These plasmids were puro resistance. The full-caPAK3 (T421E), a constitutively active mutation of PAK3, was cloned to pCDNA3.1-Hygro. H1299 cells were transfected with plasmid and selected at 400ug/ml hygromycin B.

### Real time quantitative RT-PCR

Tissues and cells were homogenized in 1 ml RNAiso^TM^ Plus lysis buffer (TAKARA). Total RNA was extracted and 2 μg RNA was transcribed into cDNA with M-MLV reverse transcriptase (Invitrogen) following the manufacturer’s instruction. SYBR Green Premix Ex Taq (Takara) was used for quantitative RT-PCR analysis. The gene-specific primers are as follows: mSmad4 sense primer, 5’-AGCCGTCCTTACCCACTGAA-3’; mSmad4 antisense primer, 5’-GGTGGTAGTGCTGTTATGATGGT-3’; hSmad4 sense primer, 5’-GCTGCTGGAATTGGTGTTGATG-3’; hSmad4 antisense primer, 5’-AGGTGTTTCTTTGATGCTCTGTCT-3’; mPAK3 sense primer, 5’-CAAAGAAACGGTCAACAACCAG-3’; mPAK3 antisense primer, 5’-AGCTGTTTTGGTATTCGACTGAT-3’; hPAK3 sense primer, 5’-AGCAATGGGCACGATTACTCC-3’; hPAK3 antisense primer, 5’-GGGCTGCTATGTATCCATGTG-3’; miR-495 sense primer, 5’-TCGCGCAAACAAACATGGTGCA-3’; miR-495 antisense primer, 5’-CAGTGCAGGGTCCGAGGTAT-3’; miR-543 sense primer, 5’-TCGCGCAAACAAACATGGTGCA-3’; mi-R-543 antisense primer, 5’-TCGCAAACATTCGCGGTGCA-3’. Technique replicates were used for every sample and each experiment was performed at least two times.

### Immunohistochemistry

Human sample study was approved by the independent ethics committee at the East China Normal University. All clinical samples were devoid of personal information. Lung cancer tumors, lung cancer metastasis tumors, normal lung samples or mouse lung cancer metastasis tissues were fixed with 4% paraformaldehyde (Beijing dingguo changsheng Biotecnology co. LTD) for 12h, and transferred into gradient ethanol, rolled, processed and embedded into paraffin. 4 μm sections were cut on a microtome (Leica, Germany). Mounted sections on to charged slides and dried 2 hours at 62℃. Blocked each sample with 100-400 μl blocking solution (NeoBioscience, ENS004.120; Boster, SA1053) for 30 minutes at room temperature to prevent non-specific binding of the antibodies. Added primary antibodies in indicated dilutions to each samples and incubated overnight at 4 degree overnight in a humidified chamber. Smad4 (Santa Cruz sc-7966, 1:200), PAK3 (Santa Cruz sc-1871, 1:100), p-c-Jun (CST #2361, 1:100), p-JNK (CST #4668, 1:50), Ras (G12D) (CST #14429, 1:50), P53 (Santa Cruz sc-6243, 1:50), TTF1 (Abcam ab76013, 1:200). Covered sections with detection reagent for 20 minutes at RT. Then stained with DAB for 1-5 minutes at RT. To evaluate the staining intensity in IHC, the white-view pictures on the slides were taken under the same condition. We utilized Image-Pro to select cells with the positive staining by calculating the grey values. The combination of size ratio between the grey areas and the whole field as well as staining intensity were calculated, and we utilized the resulting index as our definition of the staining intensity (+, +++, etc.). The negative (-) means the percentage of the positive cells with less than 25% and near background staining; the index of the positive cells between 25% and 50% with low intensity is considered as weak (+); those between 50% and 75% with intermediate staining refers to moderate staining (++); those higher than 75% or more than 50% with strongest staining represents intense staining (+++).

### Western blotting analysis

The cells or tissues were collected, washed with cold PBS buffer, and lysed using lysis buffer (RIPA buffer, 89900, Thermo Fisher) on ice for 30 minutes. Proteins were harvested from the lysates, and protein concentrations were quantified using BCA kit (Thermo Scientific #23227) following the instruction. Equal amount of protein from each sample was lysed in SDS sample buffer and resolved in 8-12% gradient SDS gels. Separated proteins were transferred to nitrocellulose membranes and immunoblotted with primary antibodies specific for PAK3 (Santa Cruz sc-1871, 1:200), c-Jun (CST #9165, 1:1000), p-c-Jun (CST #2361, 1:1000), JNK(CST #9258,1:1000), p-JNK(CST #4668, 1:1000), Smad4 (CST #46535, 1:1000), GAPDH (CST #5174 1:1000), p-MEK1(Ser298) (Abcam, ab96379), MEK1(Abcam, ab32576), E-Cadherin (CST, #3195), TWIST1 (CST, #69366), SNAIL1 (Abcam, ab216347), β-actin (MBL M177-3, 1:5000) overnight at 4℃. After incubation with a fluorescent-labelled secondary antibody (Invitrogen; Jackson Immunoresearch, 1:5,000 dilutions), specific signals for proteins were visualized by a LI-COR Odyssey Infrared Imaging System.

### Transwell assay

Transwell assays were performed in 24-well PET inserts (Millipore, 8.0-μm pore size) for cell migration. 5 ×10^4^ cells in serum-free media were plated in the upper chamber of transwell inserts (Millipore PIRP12R48) (two replicas for each sample) for 12-20 hours. The inserts were then placed into 10% serum media for indicated hours of migration as described in figures legend. Cells in the upper chambers were removed with a cotton swab, and migrated cells were fixed in 4% paraformaldehyde and stained with 0.5% crystal violet. Filters were photographed, and the total migrated cells were counted. This experiment was repeated independently three times.

### Wound healing assay

For wound healing assay, cells cultured in the 12-well plate were scratched by a small pipette tip to produce a “wound”, and monitored the “healing” after 0 hour, 6 hour, or 12 hour. The images of each well were captured, and the closure in each well were counted. Every experiment was repeated independently three times.

### F-actin staining

The cover slides were treated by 0.01% Poly-L-Lysine (Sigma) for 30min and 10μg/mL Fibronectin (Corning) for 12h, placed in a 24-well plate, and inoculated with 5×10^4^ cells per well. The cells were switched to non-serum medium for 24h, followed by 10% serum medium for 0, 15 min. The cells were washed with PBS at 37℃, fixed by 4% paraformaldehyde, permeated by 0.25% Triton X −100, stained by Rhodamine-Phalloidin (Invitrogen) and DAPI (Thermo, avoid light). The cover slips were then mounted on slides for microscopic visualization.

### Luciferase assay

pcDNA3.1-Luciferase-MCS vector was a gift from Dr. Ping Wang (TongJi University). pcDNA3.1-Luciferase reporter constructs were generated by cloning the 3’UTR of PAK3 (wild type/ mutant) at the location of MCS of pcDNA3.1-Luciferase-MCS vector. Co-transfection of WT or mutant pcDNA3.1-Luciferase reporter constructs and miRNA mimics or negative control mimics into cells. After 48 hours, removed medium and rinsed cells with PBS twice. Then used Luciferase Reporter Assay system (Promega) and record the firefly luciferase activity measurement.

### CHIP assay

PK and SPK cells were transfected with flag-PRK5-SMAD4 or flag-PRK5 vector plasmids for 36h, immersed in RPMI-1640 medium supplemented with 1% formaldehyde and protease inhibitor (Roche, USA) for 5 min. And then 125 mM glycine was added for 3 min. Cells were washed and harvested. Cells were re-suspended in sonic buffer and then sonicated until clear to result in DNA fragments of 100–500 bp in length. Wash the protein G magnetic beads (Invitrogen, 10612D) with ripa 0.3 buffer, add flag antibody (Abcom, ab49763) and rotated at 4 degrees for six hours. Then, 1ml samples were mixed with Magnetic Protein G Beads, rotated at 4 degrees overnight. Beads were washed with ripa 0.3 buffer, ripa 0 buffer, LiCl buffer, and TE buffer, respectively. Immunocomplexes were extracted from the beads with SDS elution buffer with RNase, and then added Proteinase K. Additionally, crosslinks were reversed at 65℃ for at least 6 h. Moreover, DNA fragments were purified with DNA purification kit. Q-PCR was conducted to assess the enrichment fold of immunoprecipitated DNA during the ChIP experiments. The sequences of the primers used are provided as follows: miR-495 Forward primer: 5’-ATGAGAATGGGTTTGGGTTG-3’, miR-495 Reverse primer: 5’-AGCCCTGGGTACTGTCCTCT-3’; miR-543 Forward primer: 5’-GCAGTAAACAACGCCATCCT-3’, miR-543 Reverse primer: 5’-GATGAGCAAGAACAACGAAGC-3’

### Bioinformatics analysis

We downloaded the raw data from a published paper (PMID: 11707567) and then sorted the patients who had lung cancer primary site tumor (n=123) and lung cancer metastasis tumor (n=7). We analyzed the expression of SMAD4 and PAK3 of these patients’ tumors and conducted the Oncomine analysis (https://www.oncomine.org/) on the published datasets by utilizing the cutoff criteria (P < 0.05; |log2 Fold change| > 1.5).

### Statistical analysis

Prism software (GraphPad Software) was used for statistical analyses. The intensity of the Western blot results was analyzed by densitometry using ImageJ software. Values were shown as mean ± s.e.m. Statistical significance between two samples was determined with two-tailed Student’s t-tests.

### Study approval

All mice studies were approved and used in accordance with institutional guidelines (East China Normal University). Human specimens used in this study have been approved by the East China Normal University.

## Supporting information

Sup. Data 1. Differentially expressed genes (DEGs) (spk-vs-pk-diff-pval-0.05-FC-2)

Sup. Data 2.Top enriched Human Diseases by the KEGG analysis of DEGs

Sup. Data 3.Top enriched Cellular Processes by the KEGG analysis of DEGs

Sup. Data 4.Genes both related to Cancer and Cell motility

Sup. Table 1. Summary of lung cancer metastasis in PK &SPK mice

Sup. Table 2. Summary of SMAD4 mutation in the MSK-IMPACT datasets

Sup. Table 3. Summary of Lung cancer patients have triple mutations in KRAS, TP53, and SMAD4

Sup. Table 4. Summary of Patient information (Figure 7)

## Data availability statement

The data that support the findings of this study are available from the corresponding author upon reasonable request. RNA-Seq data has been deposited into GEO database (GSE164436).

## AUTHOR CONTRIBUTIONS

Conceptualization:LT, Jian L. LL, XY, and XTL; Investigation: LT, SHS, XHT, JJX, LJ, and QWL; Resources: DS, XY, MR, JSZ, YXW, BN, LYT, JJF and KW; Bioinformatics analyses: Jian L. and FRG; Funding Acquisition and Supervision: JL, LL, XY, and XTL; Visualization and writing: JL, Jian L. LL and XTL.

## Acknowledgement

This work was supported by the National Basic Research Program of China (2015CB910402). This work was also supported in part by grants from National Natural Science Foundation of China (81672883, 81261120555, 31200878, 31071875, 81271742, 31401012 and 31730017), and Shanghai natural science foundation (17ZR1407900, 14ZR1411400).

## Supplementary Files

**Figure S1.**
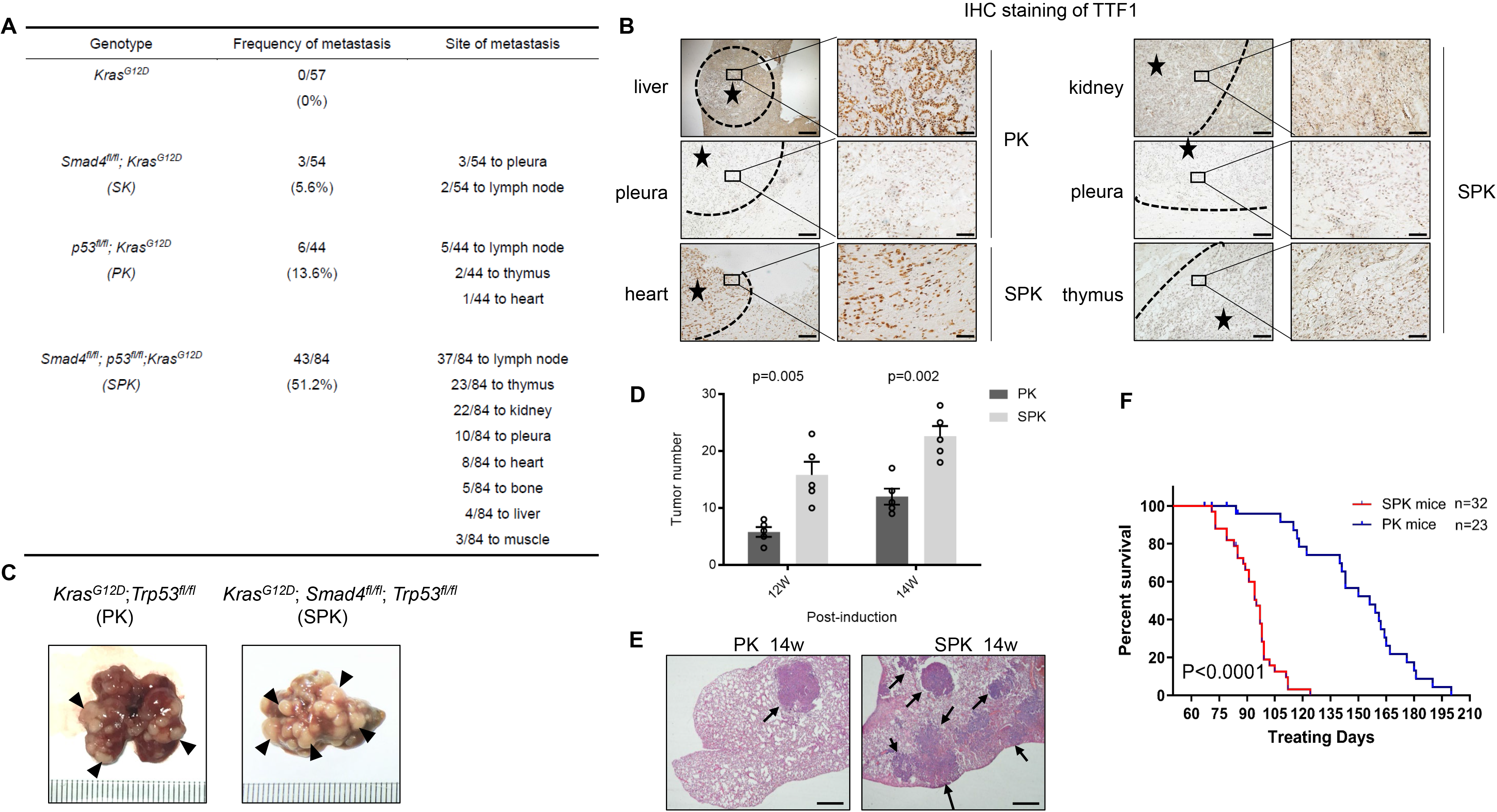
Smad4 deletion accelerates lung tumorigenesis and metastasis. **A.** Statistical analyses of lung cancer metastases frequency and the site of metastasis in *Kras^G12D^* (n=57), *Smad4^fl/fl^; Kras^G12D^* (n=54), *p53^fl/fl^; Kras^G12D^* (n=44), *Smad4^fl/fl^; p53^fl/fl^; Kras^G12D^* (n=84) mice. **B.** Representative expression of TTF1 marker in PK and SPK adenocarcinoma and metastatic tumor tissues, including liver, pleura from PK mouse and heart, kidney pleura, thymus from SPK mouse. Asterisk indicates the area of metastatic tumors. Scale bar, 100 μm (magnification, ×10), 25 μm (magnification, ×40). **C-E.** Representative pictures, H&E staining results and Statistical analyses of lung tumors of PK or SPK mouse. PK and SPK mouse took the same amount of time to treat with Ad-Cre. Scale bar, 100 μm (magnification, ×10). P<0.01, data represent means ± s.e.m.; as determined by Student’s *t* test. **F.** Survival statistics of PK and SPK mice with lung cancer. n=23 in PK and n=32 in SPK mouse groups. P<0.001. The survival range of all the mouse was between 10 and 28 weeks after the mouse infected with adeno-Cre.

**Figure S2.**
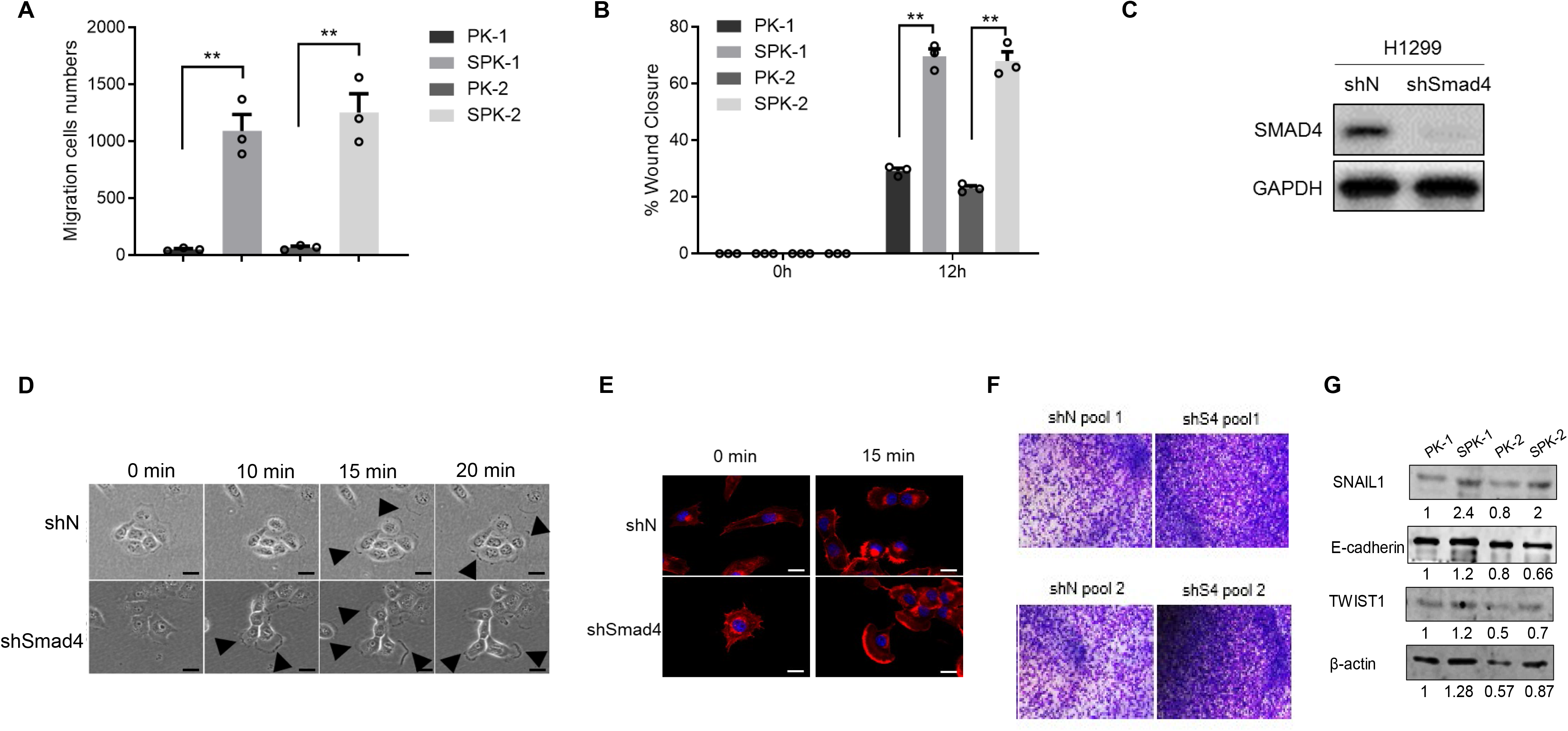
Abrogation of Smad4 promotes the lung cancer cell migration and invasion. **A.** Statistics of PK1/2 and SPK1/2 migration cells number of transwell assay(Figure 2B). Data represent means ± s.e.m. **, p<0.01, as determined by Student’s t test. **B.** Wound healing percentage of PK1/2 and SPK1/2 of Figure 2C was quantified. Data represent means ± s.e.m. **, p<0.01, as determined by Student’s t test. **C.** Detection of the expression of Smad4 in H1299 shN and H1299 shSmad4 cells. **D.** H1299 shN and H1299 shSmad4 cells were serum-starved 24 h, then stimulated with serum in Super-resolution Multiphoton Confocal Microscope and taken pictures each 30 sed. The black triangles refer to the pseudopodium. **E.** H1299 shN and H1299 shS4 cells were serum-starved 24 h before stimulation with serum for 0min and 15min at 37 °C. Cells were then fixed and stained with Rhodamine-Phalloidin (Red, F-actin) and DAPI (Blue, nucleus). The speed of cytoskeleton reconstruction, the number and length of cilia and pseudopodium, the change of cell morphology all represent the cell migration activity. The white triangles indicate cilia and pseudopodium. Scale bar, 25 μm (magnification, ×40). **F.** Invasion chamber analysis of H1299 cells, stained by 0.5% crystal violet. ShN: shRNA for control; shS4: shRNA for *SMAD4*. **G.** Western blot analysis of protein expression in PK and SPK cells.

**Figure S3.**
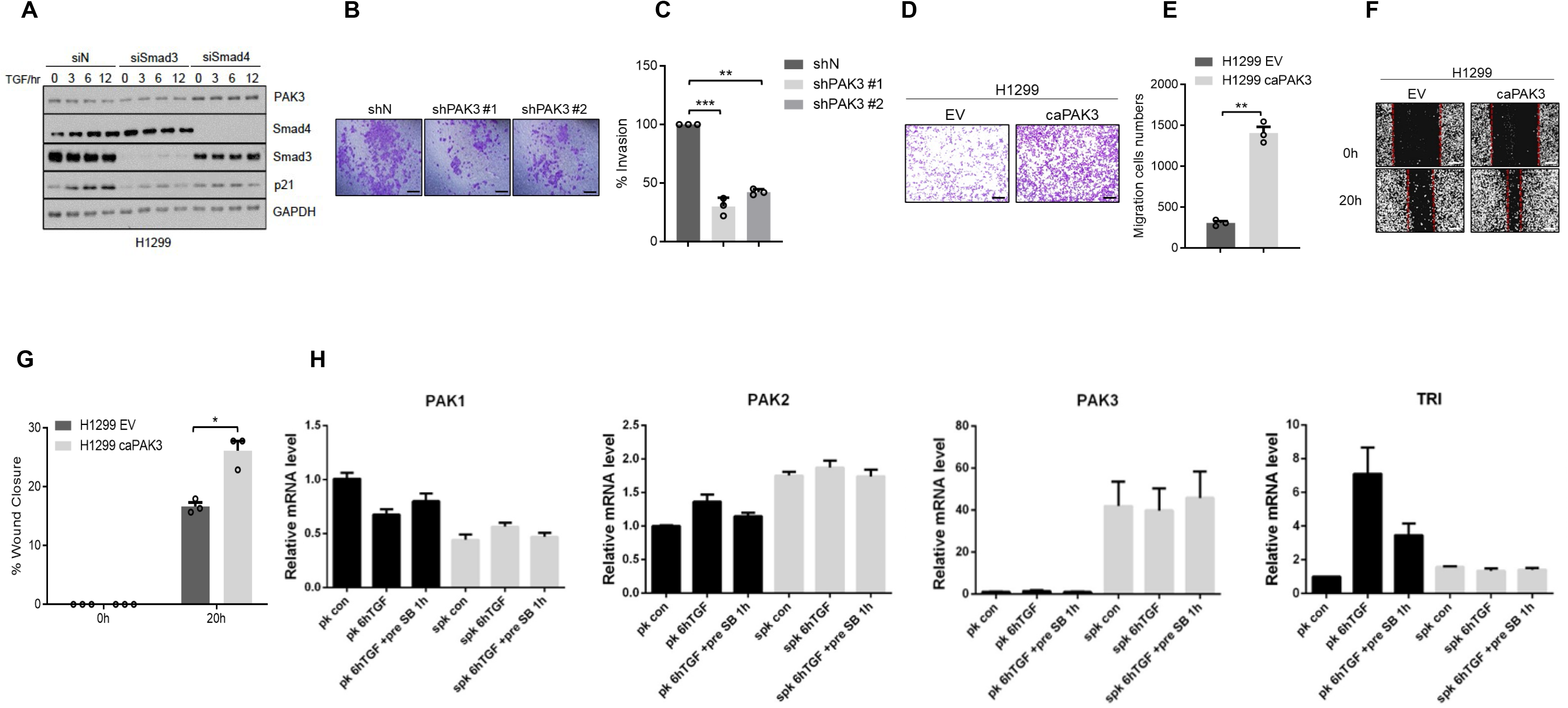
PAK3 is a downstream effector of Smad4 mediating lung cancer cell metastasis. **A.** Expression of PAK3, Smad3 and Smad4 when silencing Smad3 or Smad4 with the treatment of TGFβ in H1299 cells. **B.** The migration ability of SPK shN and shPAK3(#1/#2) cells was tested by Transwell assay. **C.** The percentage of invasive cells of Transwell assay. Data represent means ± s.e.m. **, p<0.01, as determined by Student’s t test. **D.** Over-expressed PAK3 in H1299 cells by adeno-associated virus (H1299-caPAK3). The migration ability of H1299-Vector and H1299-caPAK3 cells was compered by Transwell assay. **E.** Statistical analyses of invasion cells. Data represent means ± s.e.m. **, p<0.01, as determined by Student’s t test. **F.** The migration ability of H1299-Vector and H1299-caPAK3 cells was compered by wound healing assay. **G.** Statistical analyses of wound closure percentage. Data represent means ± s.e.m. *, p<0.05, as determined by Student’s t test. **H.** qRT-PCR analysis of gene expression in PK and SPK cells.

**Figure S4.**
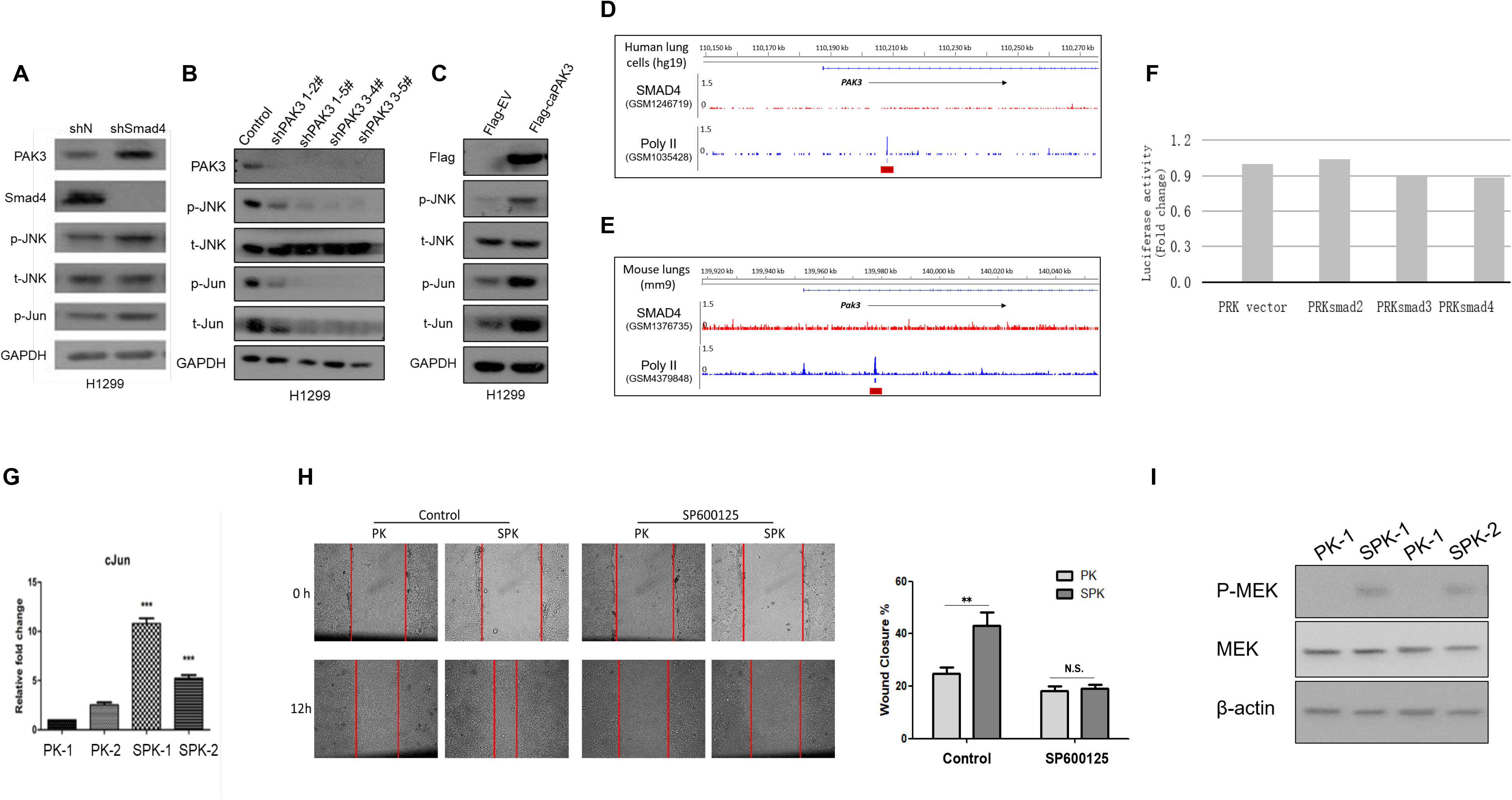
PAK3 enhances the JNK-Jun signal pathway. **A.** Expression of PAK3, JNK-Jun related factors in H1299 shN and H1299 shSmad4 cells. **B.** Expression of PAK3, JNK-Jun related factors in H1299 and several H1299 shPAK3 (inhibited expression of PAK3) cells. **C.** Expression of PAK3, JNK-Jun related factors in H1299 and H1299 Flag-caPAK3 cells. **D-E**. Analysis of the SMAD4 ChIP-Seq in human (GSM1246719) and mouse (GSM1376735) lungs in the published database. **F.** Luciferase assay on mouse Pak3 Promoter. **G.** RT-qPCR analysis of gene expression in PK and SPK cells. **H.** Migration assay of PK and SPK cells. **I.** Western blot analysis of protein expression in PK and SPK cells.

**Figure S5.**
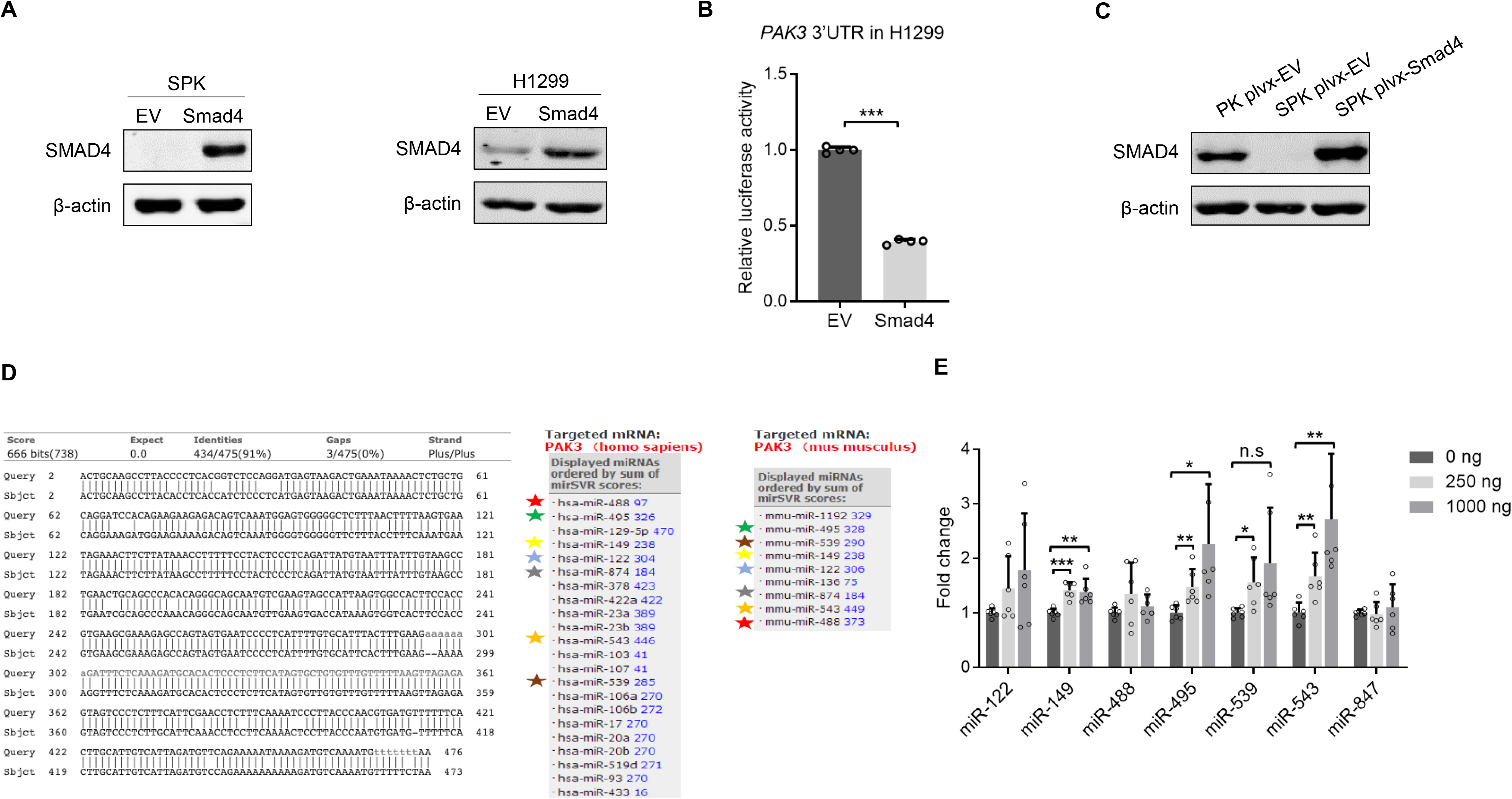
Smad4 negatively regulates PAK3 via transactivation of miR-495 and miR-543 expression. **A.** The expression of SMAD4 in normal and overexpressing SMAD4 SPK cells was detected by western blotting assay. **B.** Luciferase reporter assay showed that overexpression of Smad4 decreased PAK3 expression level in H1299 cells. The expression of SMAD4 in normal and overexpressing SMAD4 H1299 cells was detected by western blotting assay. **C.** The expression of SMAD4 in normal PK cells, normal and overexpressing SMAD4 SPK cells was detected by western blotting assay. **D.** The homology of the 3’UTR regions of murine and human PAK3 (left panel) and several PAK3 highly correlated miRNAs were evaluated (right panel). **E.** Three microRNAs, miR-495, miR-539 and miR-543, could be positively regulated by Smad4 in a dose-dependent manner in H1299 cells.

**Figure S6.**
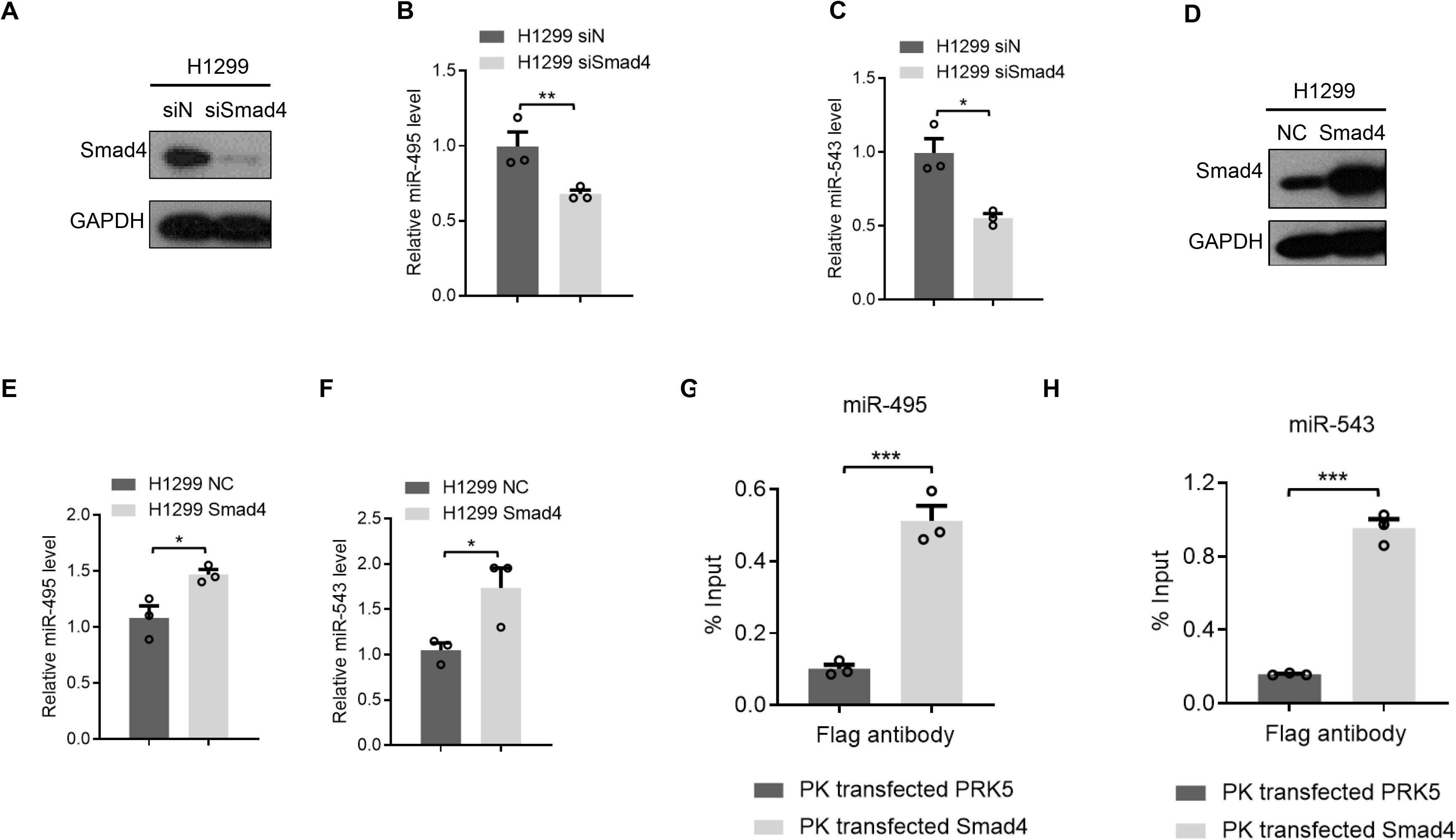
miR-495 and miR-543 were direct targets of Smad4 activation. **A.** The expression of SMAD4 in H1299 shN and H1299 shSmad4 cells was detected by western blotting assay. **B-C.** The RNA expression of miR-495 and miR-543 were detected in H1299 shN and H1299 shSmad4 cells. Data represent means ± s.e.m. *, p<0.05; **, p<0.01, as determined by Student’s t test. **D.** The expression of SMAD4 in normal and overexpressing SMAD4 H1299 cells was detected by western blotting assay. **E-F.** The RNA expression of miR-495 and miR-543 were detected in normal and overexpressing SMAD4 H1299 cells. Data represent means ± s.e.m. *, p<0.05, as determined by Student’s t test. **G-H.** The interaction between Smad4 and miR-543 in SPK cells were verified by ChIP assay. Data represent means ± s.e.m. ***p<0.001, as determined by Student’s t test.

**Figure S7.**
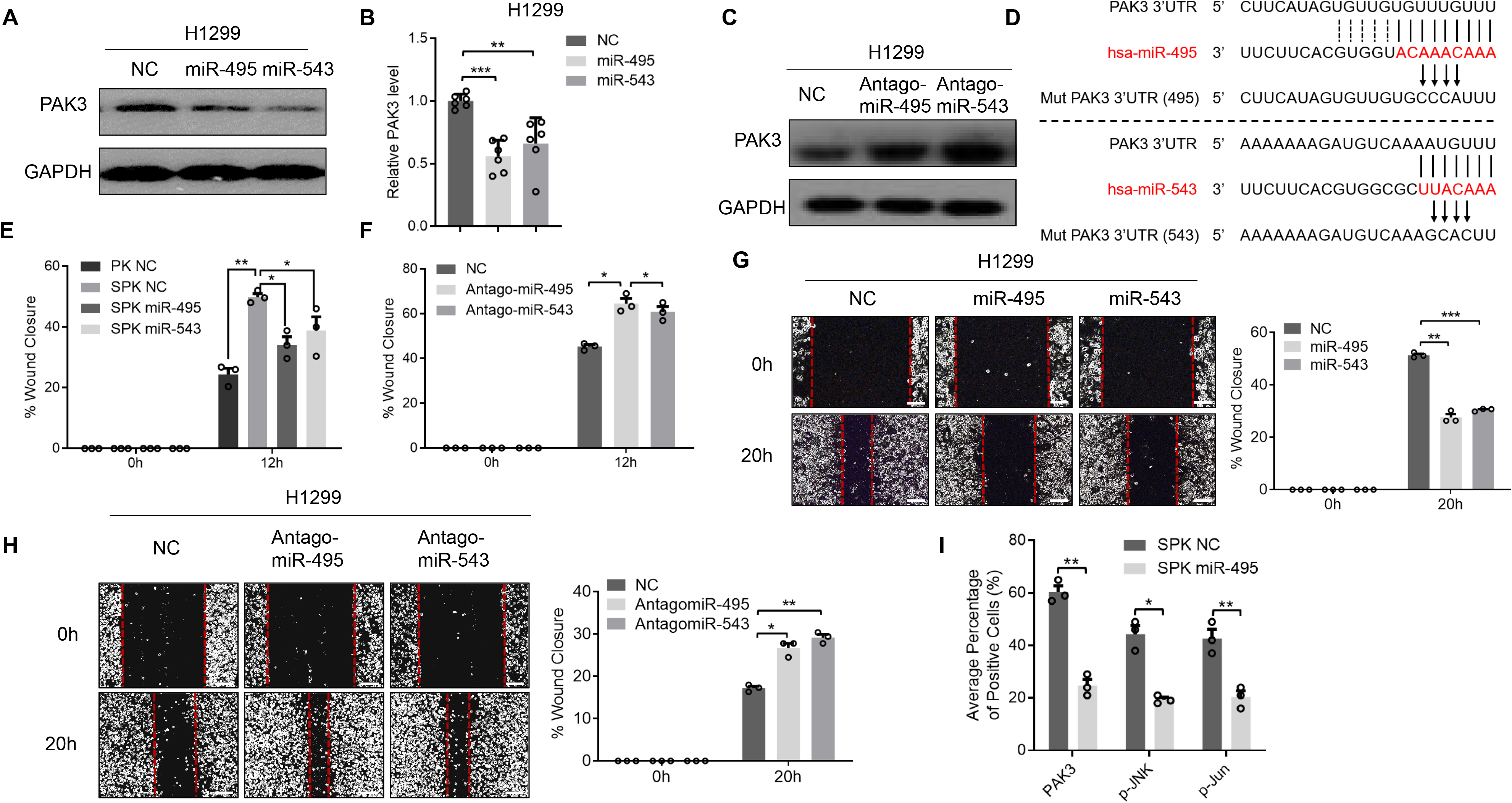
MiR-495 and miR-543 directly bind to the PAK3, UTR and attenuate metastatic potential of lung cancer cells *in vitro* and *in vivo*. **A-B.** The protein and mRNA expression level of PAK3 was tested by western blotting and qPCR in H1299 NC, H1299 miR-495 and H1299 miR-543 cells. Data represent means ± s.e.m. ***, p<0.001, as determined by Student’s t test. **C.** The protein expression level of PAK3 was detected by western blotting in H1299 NC, H1299 antagomiR-495 and H1299 antagomiR-543 cells. **D.** The sketch map of PAK3 3’UTR mutant (miR-495/miR-543) sequence. **E-F.** Statistical analyses of wound closure percentage in Figure 6F and 6G. Data represent means ± s.e.m. **, p<0.01, *, p<0.05, as determined by Student’s t test. **G-H.** The migration ability was verified by wound healing assay in H1299 cells which were treated with miR-495/543 or antogomiR-495/543 separately (left). Statistical analyses of wound closure percentage (right). Data represent means ± s.e.m. *, p<0.05; **, p<0.01; ***, p<0.001, as determined by Student’s t test. **I.** Statistics of positive cells average percentage of Fig.6H. Data represent means ± s.e.m. *, p<0.05; **, p<0.01, as determined by Student’s t test.

**Figure S8.**
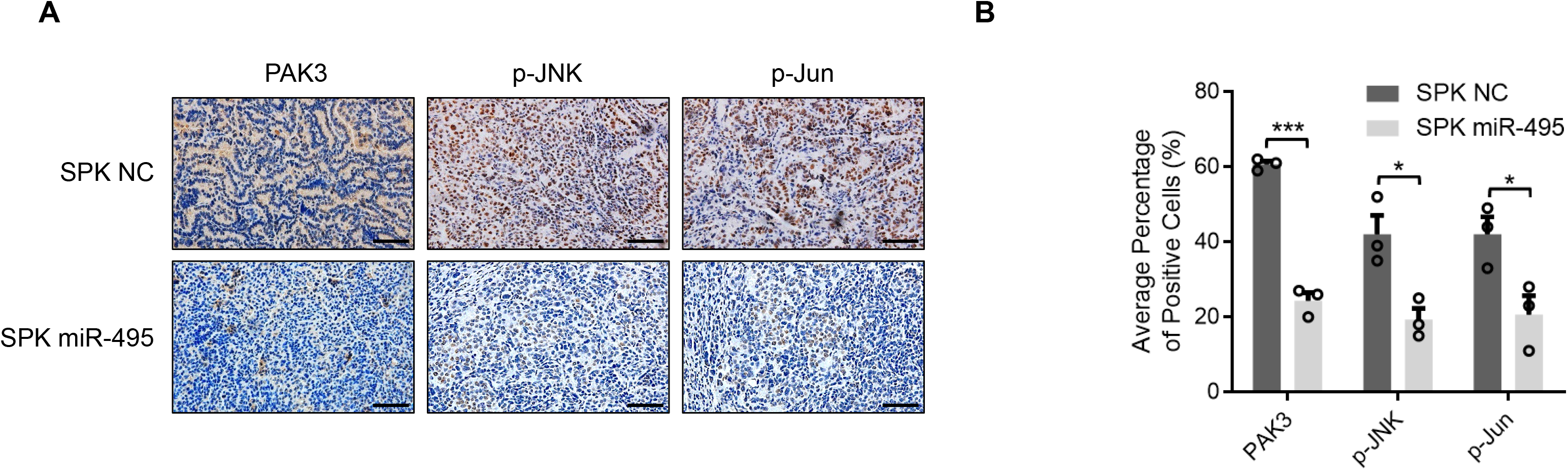
RAS (G12D) and P53 high expressed in human lung cancer metastatic tissues. **A.** The expression level of PAK3, p-JNK and p-Jun of the other parts of SPK NC and miR-495 mouse lung cancer tumors were detected by IHC staining. Scale bar, 25 μm (magnification, ×40). **B.** Statistics of positive cells average percentage of (A). Data represent means ± s.e.m. *, p<0.05; ***, p<0.001, as determined by Student’s t test.

**Figure S9.**
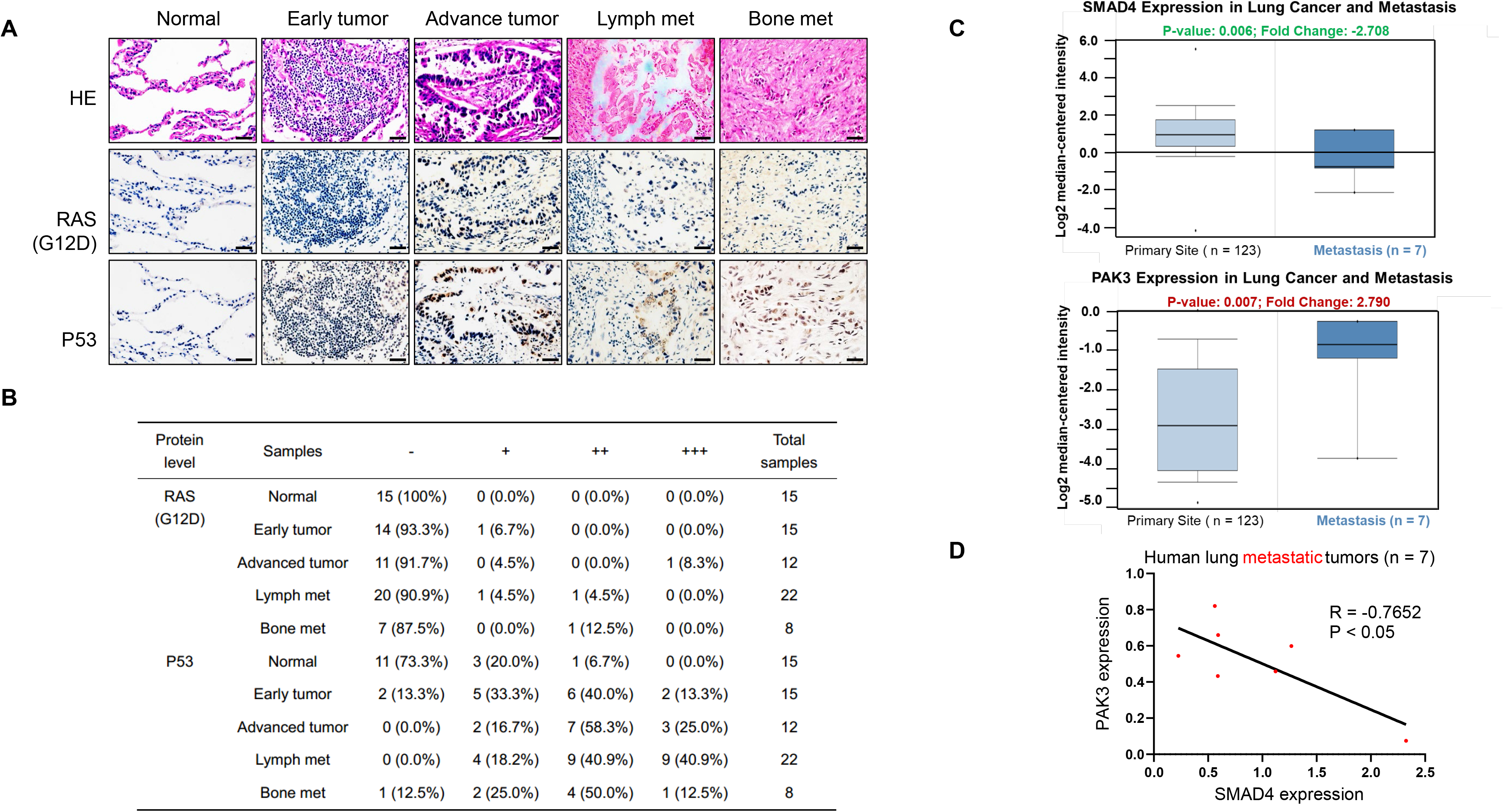
The expression of Smad4 and PAK3/JNK/Jun/RAS (G12D)/P53 in human early/advanced lung cancer tissues. **A.** Representative HE and IHC images of RAS (G12D)/P53 in human normal lung tissues, early/advanced lung cancer tissues and lung cancer lymph/bone metastatic tumor tissues. Scale bar, 25 μm (magnification, ×40). **B.** Statistics of positively stained percentages in human metastatic lung cancer and normal lung tissues (n=15); early tumor (n=15); advanced tumor (n=12); lymph nodes metastases (n=22); bone metastases (n=8). **C.** The gene expression information of 123/7 cases of lung cancer patients primary/metastasis tumor tissues from the raw data of a published paper (PMID: 11707567) were collected and analyzed the expression of SMAD4 and PAK3. **D.** Pearson correlation analysis of expression between PAK3 and SMAD4 in human lung cancer metastatic tumors(n=7). P<0.05, as determined by Student’s t test.

**Figure S10.**
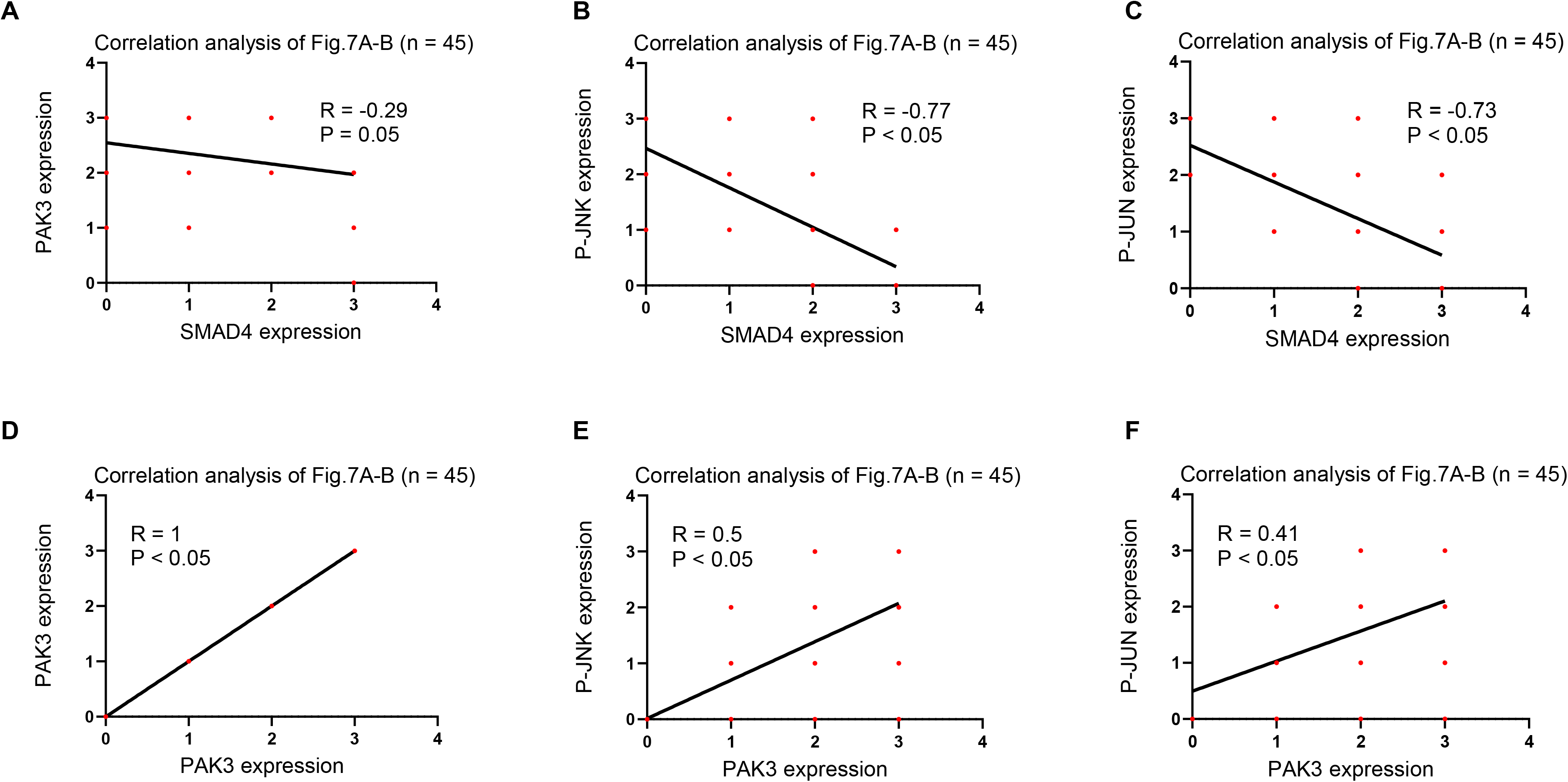
Pearson correlation analysis of Fig.7A-B. **A.** Pearson correlation analysis of expression between PAK3 and SMAD4 in human samples in Fig.7A-B. P<0.05, as determined by Student’s t test. **B.** Pearson correlation analysis of expression between p-JNK and SMAD4 in human samples in Fig.7A-B. P<0.05, as determined by Student’s t test. **C.** Pearson correlation analysis of expression between P-JUN and SMAD4 in human samples in Fig.7A-B. P<0.05, as determined by Student’s t test. **D.** Pearson correlation analysis of expression between PAK3 and PAK3 in human samples in Fig.7A-B. P<0.05, as determined by Student’s t test. **E.** Pearson correlation analysis of expression between PAK3 and P-JNK in human samples in Fig.7A-B. P<0.05, as determined by Student’s t test. **F.** Pearson correlation analysis of expression between PAK3 and P-JUN in human samples in Fig.7A-B. P<0.05, as determined by Student’s t test.

Sup. Data

Sup. Data 1. Differentially expressed genes (DEGs) (SPK cells vs PK cells).

Sup. Data 2. Top enriched human diseases by the KEGG analysis of DEGs (SPK cells vs PK cells). Sup. Data 3. Top enriched Cellular Processes by the KEGG analysis of DEGs (SPK cells vs PK cells).

Sup. Data 4. Genes both related to Cancer and Cell motility

Supplementary Tables

Sup. Table 1. Summary of lung cancer metastasis in PK &SPK mice.

Sup. Table 2. SMAD4 mutation in the MSK-IMPACT datasets.

Sup. Table 3. Summary of Lung cancer patients have triple mutations in KRAS, TP53, and SMAD4.

Sup. Table 4. Summary of Patient information.

## References

1. Siegel, R.L., Miller, K.D. & Jemal, A. Cancer statistics, 2015. CA Cancer J Clin 65, 5–29 (2015).

2. Siegel, R.L., Miller, K.D. & Jemal, A. Cancer statistics, 2016. CA Cancer J Clin 66, 7–30 (2016).

3. Siegel, R.L., Miller, K.D. & Jemal, A. Cancer Statistics, 2017. CA Cancer J Clin 67, 7–30 (2017).

4. Siegel, R.L., Miller, K.D. & Jemal, A. Cancer statistics, 2018. CA Cancer J Clin 68, 7–30 (2018).

5. Siegel, R.L., Miller, K.D. & Jemal, A. Cancer statistics, 2019. CA Cancer J Clin 69, 7–34 (2019).

6. Sharma, S.V., Bell, D.W., Settleman, J. & Haber, D.A. Epidermal growth factor receptor mutations in lung cancer. Nat Rev Cancer 7, 169–181 (2007).

7. Mills, N.E., Fishman, C.L., Rom, W.N., Dubin, N. & Jacobson, D.R. Increased prevalence of K-ras oncogene mutations in lung adenocarcinoma. Cancer Res 55, 1444–1447 (1995).

8. Reynolds, S.H. et al. Activated protooncogenes in human lung tumors from smokers. Proc Natl Acad Sci U S A 88, 1085–1089 (1991).

9. Mitsudomi, T. et al. Mutations of ras genes distinguish a subset of non-small-cell lung cancer cell lines from small-cell lung cancer cell lines. Oncogene 6, 1353–1362 (1991).

10. Takahashi, T. et al. p53: a frequent target for genetic abnormalities in lung cancer. Science 246, 491–494 (1989).

11. Farago, A.F., Snyder, E.L. & Jacks, T. SnapShot: Lung cancer models. Cell 149, 246–246 e241 (2012).

12. Gu, H., Zou, Y.R. & Rajewsky, K. Independent control of immunoglobulin switch recombination at individual switch regions evidenced through Cre-loxP-mediated gene targeting. Cell 73, 1155–1164 (1993).

13. Jonkers, J. & Berns, A. Conditional mouse models of sporadic cancer. Nat Rev Cancer 2, 251–265 (2002).

14. Tuveson, D.A. et al. Endogenous oncogenic K-ras(G12D) stimulates proliferation and widespread neoplastic and developmental defects. Cancer Cell 5, 375–387 (2004).

15. Jonkers, J. et al. Synergistic tumor suppressor activity of BRCA2 and p53 in a conditional mouse model for breast cancer. Nat Genet 29, 418–425 (2001).

16. Ji, H. et al. LKB1 modulates lung cancer differentiation and metastasis. Nature 448, 807–810 (2007).

17. Ozawa, H. et al. SMAD4 Loss Is Associated with Cetuximab Resistance and Induction of MAPK/JNK Activation in Head and Neck Cancer Cells. Clin Cancer Res 23, 5162–5175 (2017).

18. Uchida, K. et al. Somatic in vivo alterations of the JV18-1 gene at 18q21 in human lung cancers. Cancer Res 56, 5583–5585 (1996).

19. Takagi, Y. et al. Somatic alterations of the DPC4 gene in human colorectal cancers in vivo. Gastroenterology 111, 1369–1372 (1996).

20. Yamamoto, T. et al. Loss of SMAD4 Promotes Lung Metastasis of Colorectal Cancer by Accumulation of CCR1+ Tumor-Associated Neutrophils through CCL15-CCR1 Axis. Clin Cancer Res 23, 833–844 (2017).

21. Vincent, T. et al. A SNAIL1-SMAD3/4 transcriptional repressor complex promotes TGF-beta mediated epithelial-mesenchymal transition. Nat Cell Biol 11, 943–950 (2009).

22. Massague, J. TGFbeta signalling in context. Nat Rev Mol Cell Biol 13, 616–630 (2012).

23. Park, J.H., Park, B. & Park, K.K. Suppression of Hepatic Epithelial-to-Mesenchymal Transition by Melittin via Blocking of TGFbeta/Smad and MAPK-JNK Signaling Pathways. Toxins (Basel) 9 (2017).

24. Bardeesy, N. et al. Smad4 is dispensable for normal pancreas development yet critical in progression and tumor biology of pancreas cancer. Genes Dev 20, 3130–3146 (2006).

25. Kojima, K. et al. Inactivation of Smad4 accelerates Kras(G12D)-mediated pancreatic neoplasia. Cancer Res 67, 8121–8130 (2007).

26. Fisher, G.H. et al. Induction and apoptotic regression of lung adenocarcinomas by regulation of a K-Ras transgene in the presence and absence of tumor suppressor genes. Genes Dev 15, 3249–3262 (2001).

27. Jackson, E.L. et al. Analysis of lung tumor initiation and progression using conditional expression of oncogenic K-ras. Genes Dev 15, 3243–3248 (2001).

28. Insall, R.H. & Machesky, L.M. Actin dynamics at the leading edge: from simple machinery to complex networks. Dev Cell 17, 310–322 (2009).

29. Zhou, W. et al. PAK1 mediates pancreatic cancer cell migration and resistance to MET inhibition. J Pathol 234, 502–513 (2014).

30. Liu, R.X. et al. p21-activated kinase 3 is overexpressed in thymic neuroendocrine tumors (carcinoids) with ectopic ACTH syndrome and participates in cell migration. Endocrine 38, 38–47 (2010).

31. Liu, J. et al. ErbB2 Pathway Activation upon Smad4 Loss Promotes Lung Tumor Growth and Metastasis. Cell Rep 10, 1599–1613 (2015).

32. Isogaya, K. et al. A Smad3 and TTF-1/NKX2-1 complex regulates Smad4-independent gene expression. Cell Res 24, 994–1008 (2014).

33. Cancer Genome Atlas Research, N. Comprehensive molecular profiling of lung adenocarcinoma. Nature 511, 543–550 (2014).

34. Cancer Genome Atlas Research, N. Comprehensive genomic characterization of squamous cell lung cancers. Nature 489, 519–525 (2012).

35. Bhattacharjee, A. et al. Classification of human lung carcinomas by mRNA expression profiling reveals distinct adenocarcinoma subclasses. Proc Natl Acad Sci U S A 98, 13790–13795 (2001).

36. Zhao, M., Mishra, L. & Deng, C.X. The role of TGF-beta/SMAD4 signaling in cancer. Int J Biol Sci 14, 111–123 (2018).

37. Shi, Y. & Massague, J. Mechanisms of TGF-beta signaling from cell membrane to the nucleus. Cell 113, 685–700 (2003).

38. Ormanns, S. et al. The Impact of SMAD4 Loss on Outcome in Patients with Advanced Pancreatic Cancer Treated with Systemic Chemotherapy. Int J Mol Sci 18 (2017).

39. Tamura, G. et al. Allelotype of adenoma and differentiated adenocarcinoma of the stomach. J Pathol 180, 371–377 (1996).

40. Howe, J.R. et al. Mutations in the SMAD4/DPC4 gene in juvenile polyposis. Science 280, 1086–1088 (1998).

41. Kang, Y.K., Kim, W.H. & Jang, J.J. Expression of G1-S modulators (p53, p16, p27, cyclin D1, Rb) and Smad4/Dpc4 in intrahepatic cholangiocarcinoma. Hum Pathol 33, 877-883 (2002).

42. Suto, T. et al. Allelotype analysis of the PTEN, Smad4 and DCC genes in biliary tract cancer. Anticancer Res 22, 1529-1536 (2002).

43. Liu, J. et al. ERBB2 Regulates MED24 during Cancer Progression in Mice with Pten and Smad4 Deletion in the Pulmonary Epithelium. Cells 8 (2019).

44. Takaku, K. et al. Gastric and duodenal polyps in Smad4 (Dpc4) knockout mice. Cancer Res 59, 6113–6117 (1999).

45. Yang, X., Li, C., Herrera, P.L. & Deng, C.X. Generation of Smad4/Dpc4 conditional knockout mice. Genesis 32, 80–81 (2002).

46. Qiao, W. et al. Hair follicle defects and squamous cell carcinoma formation in Smad4 conditional knockout mouse skin. Oncogene 25, 207–217 (2006).

47. Xu, X. et al. Induction of intrahepatic cholangiocellular carcinoma by liver-specific disruption of Smad4 and Pten in mice. J Clin Invest 116, 1843–1852 (2006).

48. Kim, B.G. et al. Smad4 signalling in T cells is required for suppression of gastrointestinal cancer. Nature 441, 1015–1019 (2006).

49. Imielinski, M. et al. Mapping the hallmarks of lung adenocarcinoma with massively parallel sequencing. Cell 150, 1107–1120 (2012).

50. Comprehensive genomic characterization of squamous cell lung cancers. Nature 489, 519–525 (2012).

51. Losi, L., Luppi, G. & Benhattar, J. Assessment of K-ras, Smad4 and p53 gene alterations in colorectal metastases and their role in the metastatic process. Oncol Rep 12, 1221–1225 (2004).

52. Huang, D. et al. Mutations of key driver genes in colorectal cancer progression and metastasis. Cancer Metastasis Rev 37, 173–187 (2018).

53. Cordenonsi, M. et al. Links between tumor suppressors: p53 is required for TGF-beta gene responses by cooperating with Smads. Cell 113, 301–314 (2003).

54. Zehir, A. et al. Mutational landscape of metastatic cancer revealed from prospective clinical sequencing of 10,000 patients. Nature medicine 23, 703–713 (2017).

55. Leung, L. et al. Loss of canonical Smad4 signaling promotes KRAS driven malignant transformation of human pancreatic duct epithelial cells and metastasis. PLoS One 8, e84366 (2013).

56. Zhang, R., Wang, M., Sui, P., Ding, L. & Yang, Q. Upregulation of microRNA-574-3p in a human gastric cancer cell line AGS by TGF-beta1. Gene 605, 63–69 (2017).

57. Huang, S. et al. Upregulation of miR-23a approximately 27a approximately 24 decreases transforming growth factor-beta-induced tumor-suppressive activities in human hepatocellular carcinoma cells. Int J Cancer 123, 972–978 (2008).

58. Jiang, T. et al. miR-19b-3p promotes colon cancer proliferation and oxaliplatin-based chemoresistance by targeting SMAD4: validation by bioinformatics and experimental analyses. J Exp Clin Cancer Res 36, 131 (2017).

59. Li, X.B. et al. Gastric Lgr5(+) stem cells are the cellular origin of invasive intestinal-type gastric cancer in mice. Cell Res 26, 838–849 (2016).

60. Wang, Y. et al. Dysregulated Tgfbr2/ERK-Smad4/SOX2 signaling promotes lung squamous cell carcinoma formation. Cancer Res (2019).

61. Tan, X. et al. Loss of p53 attenuates the contribution of IL-6 deletion on suppressed tumor progression and extended survival in Kras-driven murine lung cancer. PLoS One 8, e80885 (2013).

